# SARS-CoV-2 envelope protein causes acute respiratory distress syndrome (ARDS)-like pathological damage and constitutes an antiviral target

**DOI:** 10.1101/2020.06.27.174953

**Authors:** Bingqing Xia, Xurui Shen, Yang He, Xiaoyan Pan, Yi Wang, Feipu Yang, Sui Fang, Yan Wu, Xiaoli Zuo, Zhuqing Xie, Xiangrui Jiang, Hao Chi, Qian Meng, Hu Zhou, Yubo Zhou, Xi Cheng, Tong Chen, Xiaoming Xin, Hualiang Jiang, Gengfu Xiao, Qiang Zhao, Lei-Ke Zhang, Jingshan Shen, Jia Li, Zhaobing Gao

## Abstract

Cytokine storm and multi-organ failure are the main causes of SARS-CoV-2-related death. However, the origin of the virus’ excessively damaging abilities remains unknown. Here we show that the SARS-CoV-2 envelope (2-E) protein alone is sufficient to cause acute respiratory distress syndrome (ARDS)-like damage *in vitro* and *in vivo*. Overexpression of 2-E protein induced rapid pyroptosis-like cell death in various susceptible cells and robust secretion of cytokines and chemokines in macrophages. Intravenous administration of purified 2-E protein into mice caused ARDS-like pathological damage in lung and spleen. Overexpressed 2-E protein formed cation channels in host cell membranes, eventually leading to membrane rupture. Newly identified channel inhibitors exhibited potent anti-SARS-CoV-2 activity and excellent protective effects against the 2-E-induced damage both *in vitro* and *in vivo*. Importantly, their channel inhibition, cell protection and antiviral activities were positively correlated with each other, supporting 2-E is a promising drug target against SARS-CoV-2.

## Introduction

Severe acute respiratory syndrome coronavirus 2 (SARS-CoV-2) has caused a worldwide pandemic. Until 26 June, 2020, there have been 9.6 million confirmed Coronavirus disease 2019 (COVID-19) cases, including more than 480 thousands deaths across the globe (Chen et al., 2020; Zhu et al., 2020). Cytokine storm and consequent acute respiratory distress syndrome (ARDS) characterized by dysfunctional immune responses and severe pulmonary injury are the main causes of SARS-CoV-2-related death(Guan et al., 2020; Mehta et al., 2020; Zhang et al., 2020). The immune system of COVID-19 patients may undergo two stages of processes after infection. In the first stage, the excessive responsive secretion of proinflammatory cytokines (IL-1β, IL-6, IL-10, TNF-α, etc.) and chemokines (CXCL10, CXCL9, CCL2, CCL3, CCL5, etc.) is readily followed by the immune system “attacking” various tissues, which cause ARDS(Xu et al., 2020). After the inflammatory outburst, the immune system in severely-illed COVID-19 patients may be programmed to an immunosuppression condition, which leads to death due to the collapse of the whole immune system (Azkur et al., 2020). Intensive studies on host-pathogen interactions have partially unveiled the immunopathogenesis of COVID-19(Cao, 2020; Grifoni et al., 2020; Peeples, 2020). Similar two other betacoronaviruses, severe acute respiratory syndrome coronavirus (SARS-CoV) and Middle East respiratory syndrome coronavirus (MERS-CoV), which could cause fatal pneumonia(de Wit et al., 2016; Zaki et al., 2012; Zhong et al., 2003), SARS-CoV-2 can also induce secretion of inflammatory cytokines and cell death by releasing pathogen-associated molecular patterns (PAMPs), such as viral RNA and damage-associated molecular patterns (DAMPs), including ATP and DNA (Merad and Martin, 2020; Tay et al., 2020). It is also very likely that SARS-CoV-2 may antagonize interferon responses in various interferon signaling pathways according to studies on SARS-CoV (Blanco-Melo et al., 2020; Merad and Martin, 2020; Tay et al., 2020). However, in addition to the dysfunctional defensive responses of host cells and the antagonism of the interferon signaling pathway by the virus, whether the SARS-CoV-2 itself possesses an offensive virulence factor that can induce cytokine storm and kills targeted cells in a direct and rapid manner remains unknown.

The SARS-CoV-2 envelope (2-E) protein is the smallest major structural protein but has been largely neglected, in contrast to other major structural proteins such as the spike (S) protein (Wrapp and Wang, 2020). Knowledge on envelope (E) protein has been limited from a few studies on the highly conserved E protein of SARS-CoV that emerged in 2003 (Zhong et al., 2003). Expression of the E protein during virus replication was thought to be essential for SARS-CoV assembly and budding (Westerbeck and Machamer, 2019). A recent study argued that the SARS-CoV-E protein does not participate in virus production but is an independent virulence factor (Nieto-Torres et al., 2014). To date, however, no study on the 2-E protein has been reported and there is no effective drug targeting the 2-E protein. Here, we showed that the 2-E protein forms a cation channel that is lethal to host cells and inhibition of the 2-E channel is new antiviral strategy.

## Results

### ● SARS-CoV-2-E induces cell death in *vitro*

The influences of 2-E on host cells were evaluated in 14 cell lines and the survival rate of each cell line were measured at 12, 24, 48 and 72 h after transient transfection of SARS-CoV-2-E plasmids (Figure 1A). Among them, Vero E6 (African green monkey kidney), 16HBE (human bronchial epithelium), A549 (human alveolar basal epithelium), HeLa (human cervical cancer) and Caco2 (human intestinal carcinoma) cells gradually died while the rest remained in a healthy state. Additionally, a conserved proline-centred β-coil-β motif mutant (SARS-CoV-2-E-4a) plasmid was also tested; this mutant allows more SARS-CoV-E protein transfer to the plasma membrane from the Golgi (Cohen et al., 2011). A relatively high expression level of 2-E was observed in the susceptible cell lines, while the rest of the cell lines displayed no expression (Figure 1A, Figure S1). The 4a mutant not only improved the 2-E expression level but also caused more severe cell death (Figure S1). Interestingly, all susceptible cells coincided with the vulnerable organs observed clinically in patients with SARS-CoV-2 infections (C. et al., 2020; Wu et al., 2020). 16HBE and Vero E6 cells were selected for further examination, and the latter was transfected with a GFP-tagged plasmid to indicate the expression of 2-E. Imaging assays and video showed that the dying cells exhibited characteristic large bubbles, similar to pyroptotic bodies from the plasma membrane, which resembled the morphological changes in Vero E6 cells infected by SARS-CoV-2 (Figure 1B and C and Video S1). Plasma membrane separation revealed that 2-E protein was translocated to membranes after expression (Figure 1D). Flow cytometry showed that the cells expressing 2-E proceeded to the Annexin V and propidium iodide double-positive stage (Figure 1E). The double-positive results represent phospha-tidylserine (PS) exposure and loss of plasma membrane integrity, which are hallmarks of pyroptosis or necrosis (Gong et al., 2017; Wang et al., 2017). Among the tested biomarkers for apoptosis, necroptosis and pyroptosis, however, levels of neither phosphorylated mixed lineage kinase domain-like protein (p-MLKL, necroptosis) nor cleaved gasdermin D (GSDMD, pyroptosis) were increased (Figure 1F, Figure S2). These data suggest that 2-E induce pyroptosis-like cell death perhaps in an atypical manner.

**Figure 1.**
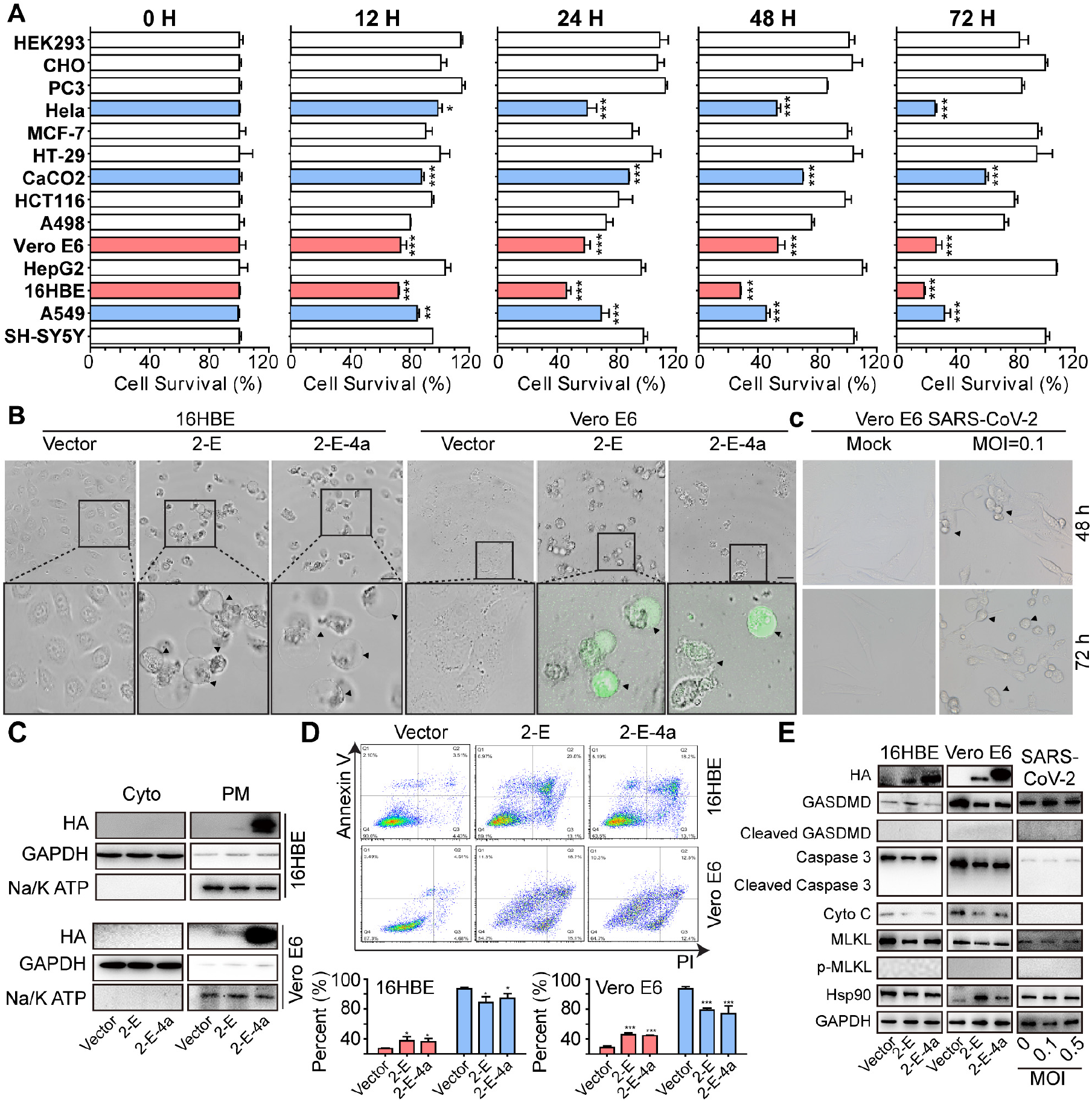
SARS-CoV-2-E expression induces cell death *in vitro*. (A) The cell viability of 14 cell lines at the indicated time after transfection with SARS-CoV-2-E plasmids. (B) SARS-CoV-2-E-induced cell death in Vero E6 cells. Phasecontrast and fluorescent images of cells. Black filled triangles indicate the pyroptosis-like bubbles (bar, 25 μm). (C) Images of Vero E6 cells infected with SARS-CoV-2 virus. (D) Subcellular fractionation analysis of SARS-CoV-2-E. Na/K ATPase is a plasma membrane marker. (E) Flow cytometry of Propidium iodide (PI) and Annexin V-FITC-stained 16HBE and Vero E6 cells. (F) Immunoblotting of cell death pathway biomarkers. All data are representative of three independent experiments. \**p* < 0.05; \*\*\**p* < 0.001; \*\*\*\**p* < 0.0001; unpaired Student’s t test. All error bars are SEM.

### ● SARS-CoV-2-E provokes robust immune responses in macrophages

Previous studies have shown that the SARS-CoV-E protein is associated with macrophage infiltration and inflammatory cytokine production (Nieto-Torres et al., 2014; Nieto-Torres et al., 2015). Three methods were used to explore the 2-E induced pathological processes in macrophages (Figure 2A). First, we transfected a mouse macrophage cell line (RAW 264.7) with SARS-CoV-2-E plasmids for 24 and 48 h. As shown in Figure 2B and Figure S3, TNF-α and IL-6 levels in cells and supernatants were significantly upregulated after transfection, as previously reported in COVID-19 patients (Merad and Martin, 2020). Second, we treated macrophages with purified 2-E protein and obtained similar enhanced cytokine levels (Figure 2C). Given that virus infection is related to innate and adaptive immune responses, it is logical to speculate that host cell damage by 2-E would lead to a secondary inflammatory cascade. To test this hypothesis, we transfected Vero E6 with 2-E plasmids, and 24 h later, the culture supernatant was collected and was used to treat RAW264.7 cells. As expected, the supernatant of impaired cells promoted cytokine expression as well, including that of IL-1RA, IL-1β and CCL5, some key markers reported for cytokine storm (Figure 2D). These data indicate that 2-E itself is sufficient to promote inflammatory responses in macrophages.

**Figure 2.**
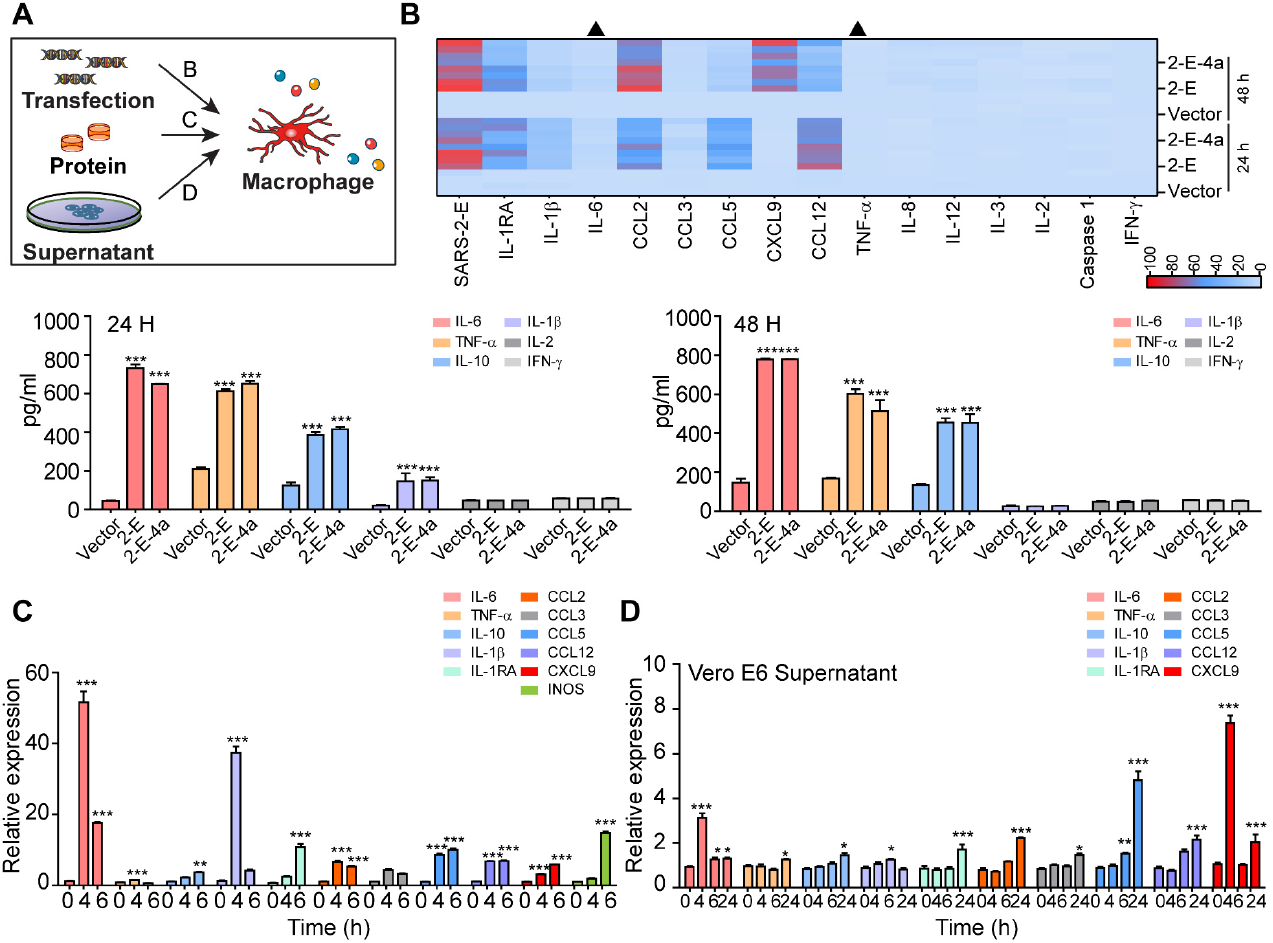
SARS-CoV-2-E provokes robust immune responses in macrophages. (A) Schematic for detecting the immune response of macrophages. (B-D) Expression of cytokines and chemokines following transfection of SARS-CoV-2-E plasmids (B), treatment with purified SARS-CoV-2-E proteins (C), and incubation with Vero E6 culture supernatant (D), measuring proteins in the supernatant via ELISA or mRNA expression via qRT-PCR. The differentially expressed genes associated with the defense response to transfection of SARS-CoV-2-E are summarized in a heatmap (B) The color code presents a linear scale. All data are representative of three independent experiments. Black triangle, time courses of the levels of IL-6 and TNF-α after transfection with 2-E plasmids (Figure S3). \**p* < 0.05; \*\*\**p* < 0.001; \*\*\*\**p* < 0.0001; unpaired Student’s t test. All error bars are SEM.

### ● SARS-CoV-2-E causes ARDS-like damage in lung and spleen *in vivo*

To explore the potential influences of 2-E protein *in vivo*, purified 2-E protein was injected intravenously for 2, 6, and 72 h in C57BL/6 mice (Figure 3A - C, Figure S4). Surprisingly, we observed severe foci of pulmonary consolidation in the lung and spleen edema at 72 h (Figure 3D), but not found in mock and bovine serum albumin (BSA) groups (Figure S5). H&E staining of lung tissue showed marked inflammatory cell infiltration, edema, pulmonary interstitial hyperemia, hemorrhage, and alveolar collapse. The observed animal phenotype was consistent with COVID-19 patients’ lungs, in which the first and most severe lesion signs appeared (Zhang et al., 2020). In addition, SARS-CoV-2 infection may trigger an overwhelming inflammatory response, which leads to injuries to multiple organs (Bao et al., 2020; Noris et al., 2020; Tay et al., 2020). We used qRT-PCR and ELISA to characterize the pathological features. As illustrated in Figure 3E and F, the expression of cytokines (IL-1RA, IL-1β, IL-6, TNF-α, etc.) and chemokines (CCL2,3, and 5, CXCL9, etc.) increased dramatically *in vivo*, along with the notably upregulated inflammatory markers, which also correlate with the previously reported cytokine storm in patients (Figure 3E and F). Our data suggest that 2-E protein alone is sufficient to drive pulmonary congestion and inflammation.

**Figure 3.**
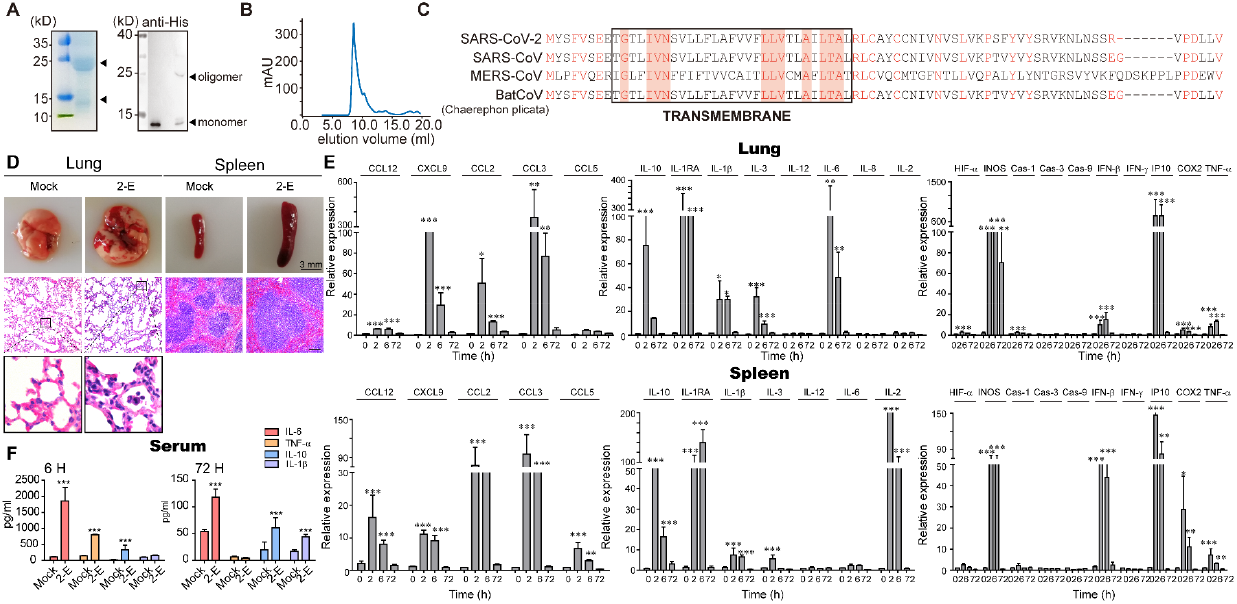
The SARS-CoV-2-E causes ARDS-like injury and inflammation in the lung and spleen *in vivo*. (A) Purification of full-length SARS-CoV-2-E protein with Ni-NTA affinity chromatography (left: 15% SDS-PAGE gel; right: Western blot probed with anti-his-tag antibodies). The four peptides of 2-E were detected by LC-MS/MS. (B) Sizeexclusion chromatogram of the affinity-purified SARS-CoV-2-E protein. Data from a Superdex 75 Increase 10/300 column are shown in blue. The elution peak probably represents SARS-CoV-2-E protein oligomers. (C) Sequence alignment of four pandemic coronaviruses (SARS-CoV-2 and SARS-CoV, MERS-CoV and BatCoV). Transmembrane residues are highlighted in the black frame. (D) Gross pathology and histopathology of lungs and spleen from control mice (Tris-buffered saline, TBS) and model mice (2-E proteins). (E) qRT-PCR analysis of lung and spleen tissues after injection of 2-E proteins. (F) Serum cytokine levels 6 h and 72 h after treatment. \**p* < 0.05; \*\*\**p* < 0.001; \*\*\*\**p* < 0.0001; unpaired Student’s t test. All error bars are SEM.

### ● SARS-CoV-2-E forms cation channels

It has been reported that the executer of pyroptosis (GSDMD) and necroptosis (p-MLKL) destroy the membrane integrity by either forming pores or channels (Ding et al., 2016; Xia et al., 2016). In addition, the E protein of SARS-CoV was found to be able to form pores with ion channel activity(Liao et al., 2006; Pervushin et al., 2009; Verdiá-Báguena et al., 2013). A planar lipid bilayer recording technique was used to assess potential electric signals mediated by 2-E. Typical single-channel currents were captured when the protein was reconstituted in asymmetric 50:500 mM KCl solutions and 3:2 PC/PS lipid membranes at applied potentials of −100 mV to +100 mV (Figure 4A). The measured reversal potential (59.25 mV) was very close to the theoretical equilibrium potential of K^+^, indicating that the 2-E channels were permeable to K^+^ but not to Cl^−^. By examining the reversal potentials in solutions containing varied concentrations of K^+^, Na^+^ and Cl^−^, we found the 2-E channels were also permeable to Na^+^ in the presence of K^+^ and Cl^−^ (Figure 4A and B). In addition, 2-E channels exhibited a much lower permeability to divalent Ca^2+^ and Mg^2+^ in the presence of K^+^ (Figure 4 A and E). The permeability rank ((*P*_Na_^+^≈*P*_K_^+^) / (*P*_Ca_^+^≈*P*_Mg_^2+^)≈3.0) suggested that 2-E channel possesses a distinct selectivity pattern from classic voltage dependent channels. The pH value on the lumen side of organelles may change after coronavirus infection (Westerbeck and Machamer, 2019). We found that both the amplitude and voltage dependence of 2-E channel-mediated currents gradually increased when pH decreased. At pH 10, the 2-E channel almost completely closed (Figure 4C and D). Collectively, our data support that 2-E forms a type of pH sensitive cation channel.

**Figure 4.**
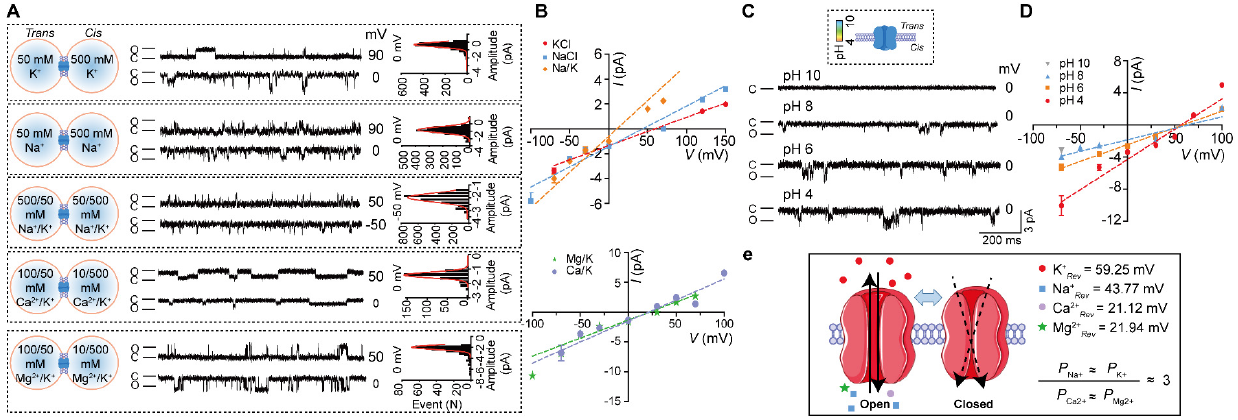
SARS-CoV-2-E forms cation channels. (A) Single-channel current recordings of 2-E after reconstitution in lipid bilayers with PC/PS = 3:2 lipids at the indicated potentials and in the indicated solutions. Protein (5-50 ng/ml) was added to the *cis* side. All-point current histograms for left the trace (0, 0, −50,0,0 mV). (B) *I-V* curves of 2-E proteins in different solutions as in the left panel (n ≥ 3). (“C” means Closed; “O” means Open) (C) Representative single-channel current recordings of 2-E in the presence of various concentrations of protons. *I-V* plot are shown in pane D (n ≥ 3). (E) Schematic diagram of 2-E channel open and close states.

### ● Newly identified channel inhibitors exhibit protective effects against 2-E-induced damage and anti-SARS-CoV-2 activity

We sought to identify inhibitors for 2-E channels using the planar lipid bilayer recording technique. Tetraethylammonium (TEA, 5 mM), Tetrodotoxin (TTX, 100 μM) and Nifedipine (100 μM), three classic inhibitors of potassium, sodium and calcium channels, respectively, showed negligible effects on 2-E channels. Although 5-(N,N-hexamethylene)-amiloride (HMA) and amantadine, two reported inhibitors of proton pumps and SARS-CoV-E channels, exhibited inhibitory activity on 2-E channels at 50 μM or greater, further investigation into these compounds was hindered by their strong cytotoxicity. BE12 (Berbamine), a type of bisbenzylisoquinoline alkaloid isolated from *Berberis*, drew our attention after screening an in-house compound collection. Although BE12 was a weak inhibitor of 2-E channels, with an IC_50_ of 111.50 μM, it exhibited minor cytotoxicity, which allowed us to further test its cell protection activity against 2-E induced cell death and anti-SARS-CoV-2 activity using Vero E6 cells. Encouraged by the promising cell protection and antiviral activity of BE12, four more channel inhibitors (BE30~33) were designed, synthesized and evaluated individually. Among them, BE33 exhibited exceptional antiviral activity against SARS-CoV-2 infection, with an IC_50_ of 0.94 μM and negligible cytotoxicity (Figure 5A-D). In addition, we treated the 2-E injected mice with BE-33 and found that BE-33 (0.50 mg/Kg) significantly reduced cytokine secretion (Figure 5E). In addition, it is noteworthy that the channel inhibition activity, the cell protection activity against 2-E-induced cell death and the antiviral activity of this class of compounds are positively correlated with each other (Figure 5F), indicating the importance of the channel activity of 2-E.

**Figure 5.**
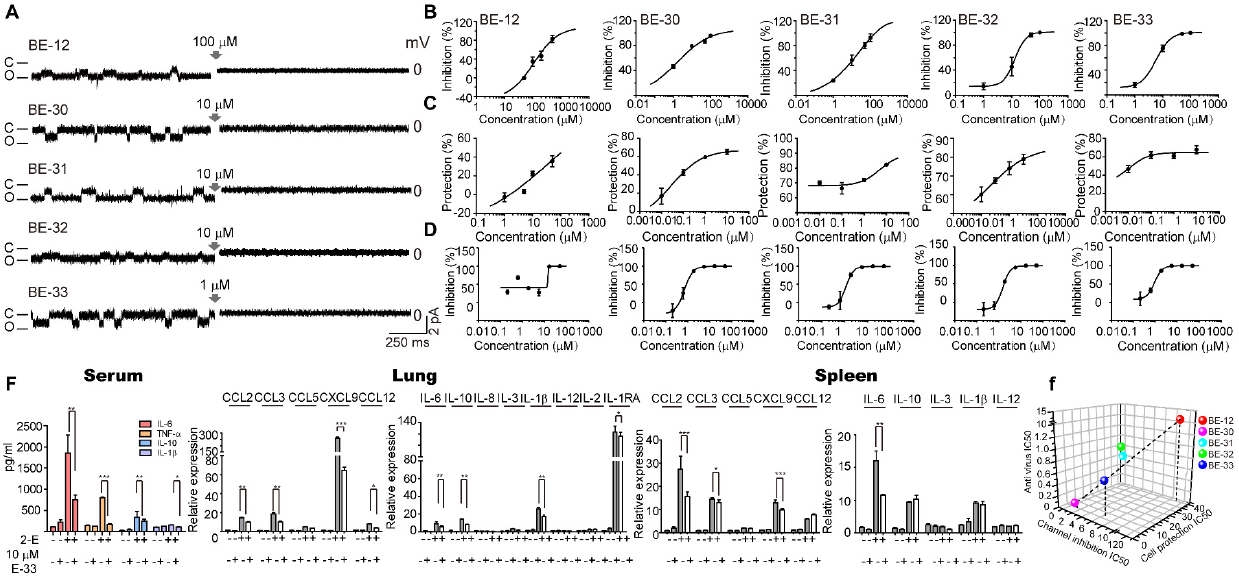
Newly identified channel inhibitors exhibit protective effects against 2-E-induced damage and anti-SARS-CoV-2 activity. (A) Representative single-channel traces after exposure to the indicated compounds at different concentrations. Once ion channel conductance was detected, compounds were added to the *trans* chamber while stirring to facilitate binding of the compound to the channel. The gray arrow indicates the application of compounds (n ≥ 3). (B-D) Doseresponse curves of the indicated inhibitors on channel activity (B) 2-E-induced cell death (C) and SARS-CoV-2 infection (D) in Vero E6 cells. (E) Expression of cytokines and chemokines in the lung and spleen after treatment with 0.50 mg/Kg BE33. (F) The correlation among the IC_50_S for channel inhibition, cell protection and antiviral activity.

## Discussion

The lack of efficient antiviral drugs for SARS-CoV-2 has prompted the urgent need for developing new therapeutic development for COVID-19. A better understanding of which viral proteins are major risk factors for organ and immune system damage is crucial to develop rationale-based clinical therapeutic strategies. Our study supports that the 2-E channel acts as an “offensive” virulence factor carried by SARS-CoV-2, leading to the robust inflammatory responses and rapid cell death. 2-E channels may act either by themselves or via other proteins to increase ion fluxes across the membranes, which changes the ion homeostasis and eventually results in membrane rupture (Feltham and Vince, 2018). In contrast to exocytosis, a common and mild mechanism for virus release, a large amount of SARS-CoV-2, 2-E and other DAMPs can be released simultaneously because of the sudden cell death induced by 2-E channels. Coincidently, COVID-19 patients’ lungs showed evident desquamation of pneumocytes and hyaline membrane formation, some COVID-19 patients may worsen rapidly to sudden stroke from mild symptoms.

It has been demonstrated that depletion of intracellular potassium will promote robust secretion of cytokines and aggravate the MLKL channel-mediated necroptosis and GSDMD pore-mediated pyroptosis (Conos et al., 2017; Ding et al., 2016). The disease severity of COVID-19 patients is linked to electrolyte imbalance, including reduced serum concentrations of potassium, sodium and calcium (Banerjee et al., 2018; Lippi and South, 2020). Identification of 2-E as a new type of potassium permeable channels may therefore unravel the pathogenic mechanisms underlying hypokalaemia exacerbating ARDS, which is a common complication in COVID-19 patients (Louhaichi et al., 2020). In addition to 2-E channels, multiple proteins exhibiting channel activity have been also identified in other highly pathogenic respiratory viruses, including SARS-CoV, MERS-CoV, and Influenza A (Cady et al., 2010; Nieto-Torres et al., 2014; Surya et al., 2015). Although how these channels modulate the ion homeostasis across the virus membrane remains unclear, it is generally believed that these channels are necessary for the virus production and maturation (Castano-Rodriguez et al., 2018). The requirement of the channels in a large number of pathogen-host interactions makes the virus channel research a field of growing interest.

One limitation of our study is the lack of evidence in the context of SARS-CoV-2 infection *in vivo*, which can technically be achieved by generating of SARS-CoV-2 without 2-E using reverse genetics. However, the 2-E deleted SARS-CoV-2 may replicate more effectively and thus this experiment was not conducted. Although the *in vivo* antiviral activity of the newly identified channel inhibitors needs further studies, their potent antiviral activity *in vitro* and excellent protection effects against ARDS-like damage *in vivo* shed light on the drug development of 2-E channel inhibitors. Given that 2-E can function as ion channels in the viral membranes, similar to how they function in host cells, we propose that 2-E channels may represent a new class of dual-function targets against SARS-CoV-2.

## Supporting information

Supplemental Vedio 1

## Acknowledgements

We thank Prof. Eric Xu for linguistic assistance during the preparation of this manuscript. We are grateful to the National Science Fund of Distinguished Young Scholars (81825021), Fund of Youth Innovation Promotion Association (2019285), the National Natural Science Foundation of China (81773707, 31700732), the Strategic Leading Science and Technology Projects of Chinese Academy of Sciences (XDA12050308), Fund of National Science and Technology Major Project (2018ZX09711002-002-006) and the Hubei Science and Technology Project (2020FCA003) for financial support.

## Author contributions

Z. G. and J. L. conceived the project. Z. G., J. L., B. X., J. S. and L.-K. Z. designed the experiments; Y. W. and B. X. performed the electrophysiological recordings; X. S., S. F. and B. X. carried out the cell-based assays; P. X. and Y. Wu. carried out the virus assays in vitro; S. F., X. S., B. X., Y. W., X. Z. and H. C. carried out the animal experiments; S. F., B. X., Y. W., Z. X., H. C., X. X., T. C. and Q. Z. purified the proteins; Q. M. and H. Z. performed the LC-MS/MS analysis; X. C. and H. J. analyzed the sequence; all authors analyzed and discussed the data. Z. G., B. X., Y. H., and L.-K. Z. wrote the manuscript.

---

**Table 1.**
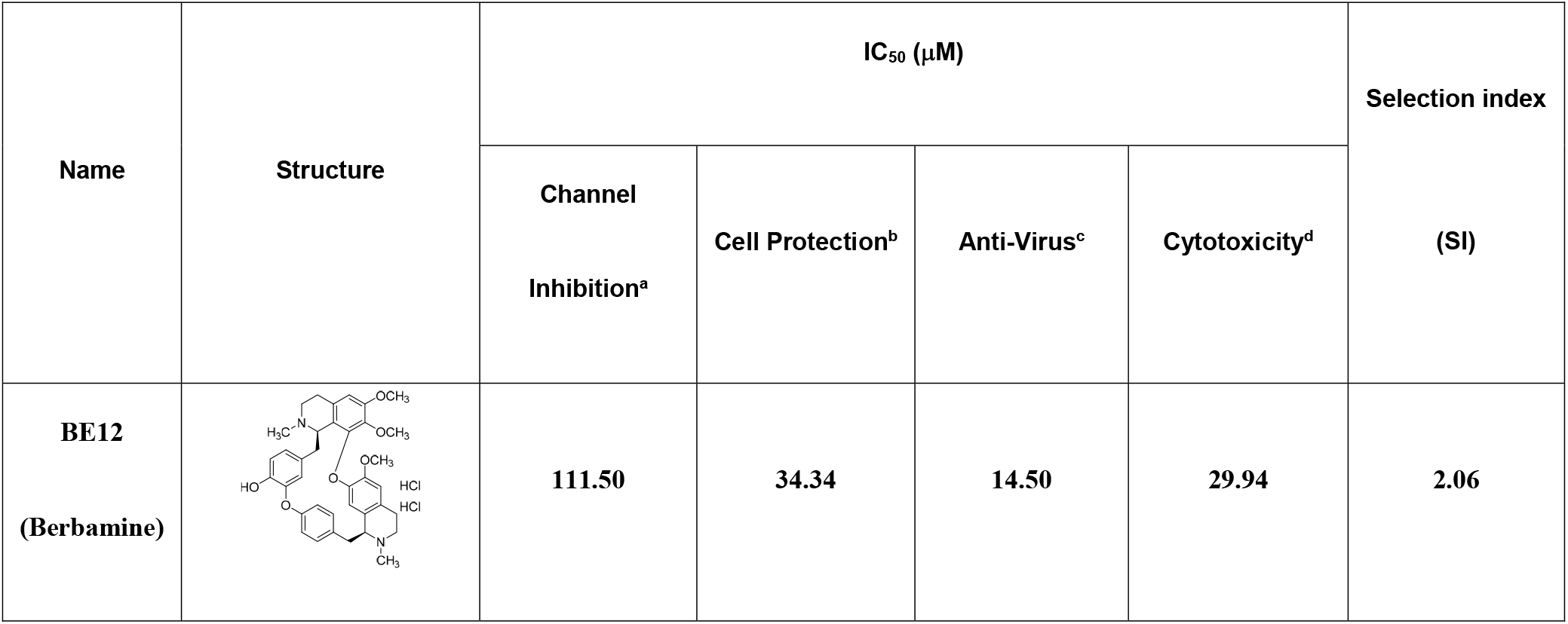
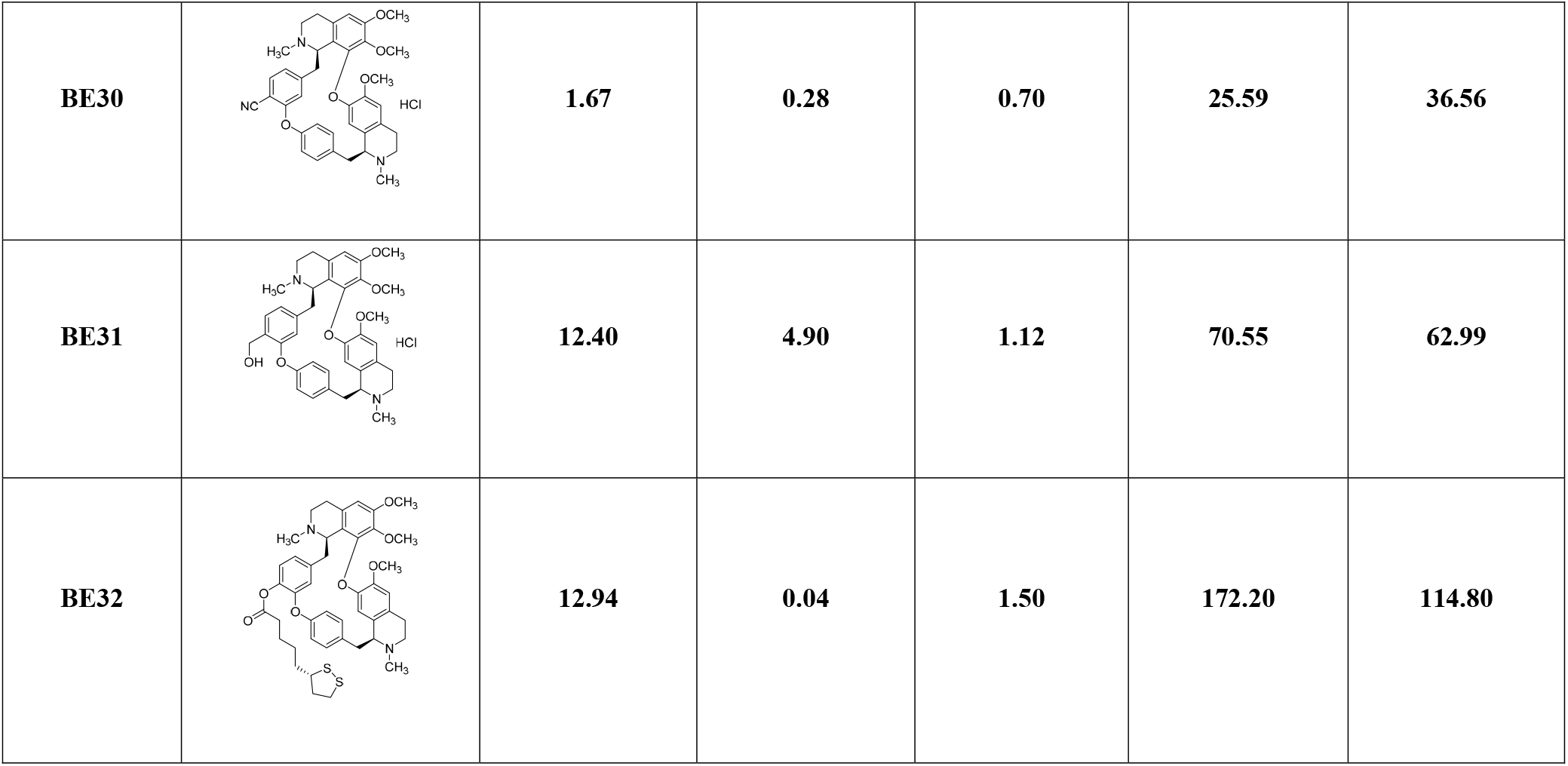
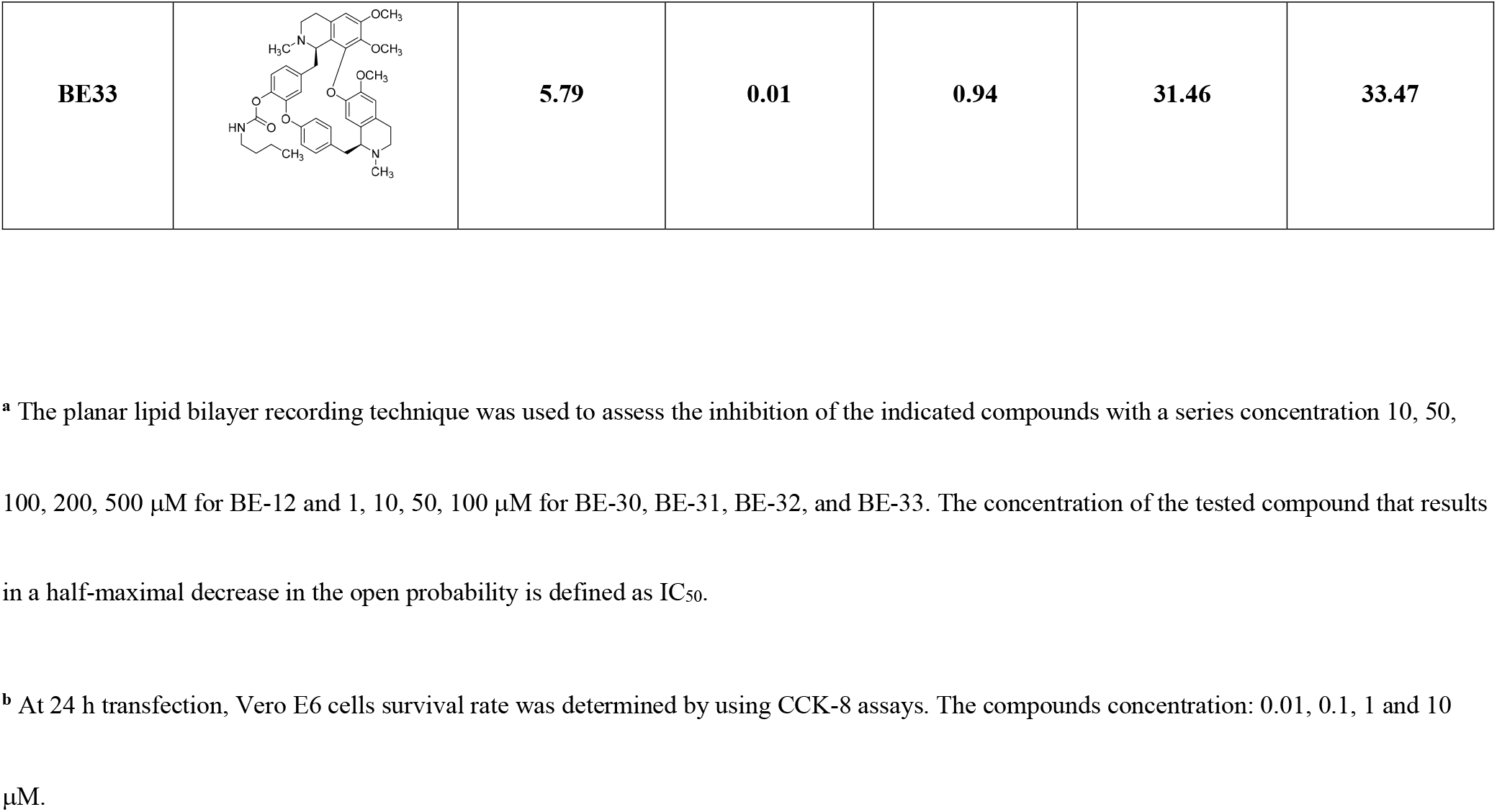
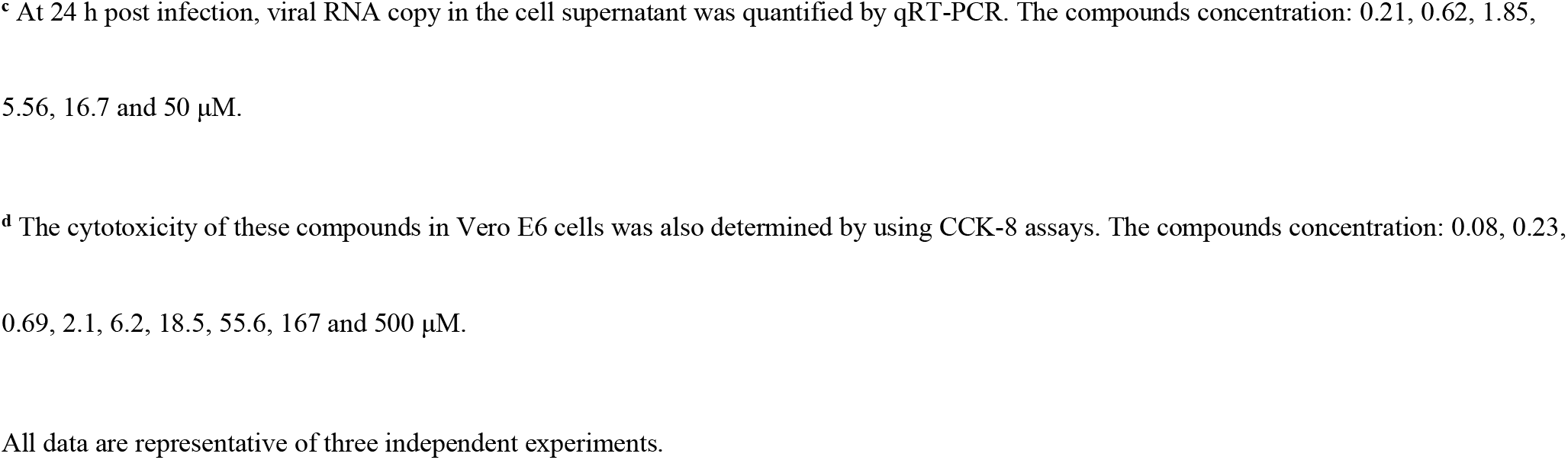
Activity of newly identified 2-E channel inhibitors *in vitro*.

| Name | Structure | IC<sub>50</sub> (μM) |  |  |  | Selection index<br>(SI) |
| --- | --- | --- | --- | --- | --- | --- |
|  |  | Channel<br>Inhibition<sup>a</sup> | Cell Protection<sup>b</sup> | Anti-Virus<sup>c</sup> | Cytotoxicity<sup>d</sup> |  |
| BE12<br>(Berbamine) |  | 111.50 | 34.34 | 14.50 | 29.94 | 2.06 |
| **BE30** |  | **1.67** | **0.28** | **0.70** | **25.59** | **36.56** |
| **BE31** |  | **12.40** | **4.90** | **1.12** | **70.55** | **62.99** |
| **BE32** |  | **12.94** | **0.04** | **1.50** | **172.20** | **114.80** |
| **BE33** |  | **5.79** | **0.01** | **0.94** | **31.46** | **33.47** |

<sup>a</sup> The planar lipid bilayer recording technique was used to assess the inhibition of the indicated compounds with a series concentration 10, 50, 100, 200, 500 μM for BE-12 and 1, 10, 50, 100 μM for BE-30, BE-31, BE-32, and BE-33. The concentration of the tested compound that results in a half-maximal decrease in the open probability is defined as IC<sub>50</sub>.
<sup>b</sup> At 24 h transfection, Vero E6 cells survival rate was determined by using CCK-8 assays. The compounds concentration: 0.01, 0.1, 1 and 10 μM.
<sup>c</sup> At 24 h post infection, viral RNA copy in the cell supernatant was quantified by qRT-PCR. The compounds concentration: 0.21, 0.62, 1.85, 5.56, 16.7 and 50 μM.
<sup>d</sup> The cytotoxicity of these compounds in Vero E6 cells was also determined by using CCK-8 assays. The compounds concentration: 0.08, 0.23, 0.69, 2.1, 6.2, 18.5, 55.6, 167 and 500 μM.
All data are representative of three independent experiments.

## STAR Methods

### Plasmids and mutagenesis

Wild type and mutant (2-E-4a: FYVY56/57/58/59AAAA) SARS-CoV-2-E sequences were synthesized by the Beijing Genomics Institute (BGI, China). Vector pET28a was used for protein purification; vector pcDNA5 was used for cell survival assay and cell imaging.

### Cell culture and treatment

HEK293, MCF-7, Caco2, Vero E6, HeLa, HepG2, SH-SY5Y cells were grown in 90% DMEM basal medium (Gibco, USA) supplemented with 10% fetal bovine serum (Gibco, USA), 2 mM L-glutamine and 100 units /mL penicillin/streptomycin (Gibco, USA). 1% NEAA (Gibco, USA) was added in above medium for A498 cells culture. Besides, HCT116 and HT-29 cells were grown in McCoy’s 5A basal medium (Gibco, USA), PC3 and A549 cells were grown in RPMI-1640 basal medium (Hyclone, USA) and CHO cells were grown in DMEM/F-12 basal medium (Gibco, USA) supplemented as above. 16HBE cells were grown in KM (ScienCell, USA) medium. For cell viability assay and western blot, cells were transfected with 3200 ng or 200 ng of SARS-CoV-2-E and SARS-CoV-2-E-4a plasmids for 6-well or 96-well by using Lipo3000 transfection reagent according to the company’s instructions (Thermo Fisher, USA). For cytotoxicity assay, Vero E6 cells were seeded in 96-well plates with 15,000 cells per well overnight. Compounds were diluted with medium to appropriate concentrations, a volume of 100 μL was added to each well and then incubated for 24 h.

### Cell viability and Cytotoxicity assay

Cell viability and Cytotoxicity were measured using the CCK8 kit (40203ES60, Yishen, China). Assays were performed according to the manufacturer’s instructions. Measure the absorbance at 450 nm with Thermo Scientific Microplate Reader (Thermo Fisher Scientific Inc., USA).

### Isolation of membrane proteins

Mem-PER™ Plus Membrane Protein Extraction Kit (89842, Thermo Fisher Scientific Inc., USA) was used to extract cytosolic and membrane proteins. Briefly, cells were washed with ice-cold PBS before the cells were harvested and permeabilized in a Permeabilization Buffer. Cytosolic and crude membrane fractions were separated by centrifugation at 3,00 × g for 5 mins. The supernatants were taken as cytosolic proteins. The pellets were washed with PBS and then solubilized to collect membrane proteins. Centrifuge tubes at 16,000 × g for 15 mins at 4°C. Transfer supernatant containing solubilized membrane and membrane-associated proteins to a new tube. Blots were probed with antibodies against HA-Tag (C29F4) (3724, CST, USA), GSDMD (L60) (93709, CST, USA), LC3A/B (4108, CST, USA), Caspase-3 (14220, CST, USA), Cytochrome c (D18C7) (11940, CST, USA), MLKL (D216N) (14993, CST, USA), Phospho-MLKL (Ser358) (D6H3V) (91689, CST, USA), HSP90 (4874, CST, USA), GAPDH (30201ES20, Yishen, China), Na/K-ATPase (3010, CST, USA).

### Flow cytometry of cell death

After treatment, cells were detached, collected by centrifugation and resuspended in 1× binding buffer containing 100 μg/mL propidium iodide (PI) and Annexin V-FITC (1:20, V13242, Life) and incubated at room temperature for 15 min in the dark. Subsequently, added 400 μL of 1× binding buffer and kept on ice. Cells were analyzed by flow cytometry using BD FACSCALIBUR 4 (BD COMPANY, USA). The percentages of differently labeled cells were calculated by FlowJo 7.6.

### Imaging

Two ways were used to examine cell death morphology. The one, cells were seeded as 2×10^5^ cells per well in the 6-well plates (30720113, Thermo Fisher Scientific Inc., USA). After transfection for 24 h, cells were captured using Leica TCS-SP8 STED system (Leica Microsystems, DE) with a 40× phase difference objective. All image data shown are representative of at least three randomly selected fields. Another, Vero E6 cells were seeded as 1×10^4^ cells per well in 24-well plate overnight. Next day, the culture medium was replaced with 2% FBS DMEM. Then SARS-CoV-2 viruses were added at MOI = 0.1, no infection was made as Mock. At 48 and 72 h, pictures of three visions were captured from each well by a fluorescence microscope (Olympus, Japan) at bright field.

### ELISA

Cell culture supernatants and serum were assayed for mouse IL-6 (VAL604, R&D Minnesota, USA), mouse TNF-α (VAL609, R&D Minnesota, USA), mouse IL-10 (VAL605, R&D Minnesota, USA), mouse IL-1β (VAL601, R&D Minnesota, USA), mouse IFN-γ (VAL 607, R&D Minnesota, USA), mouse IL-2 (abs520002, Absin, China) according to the manufacturer’s instructions.

### Quantitative Real-time PCR (qRT-PCR) Analysis

Total RNAs were extracted from cells and tissues using Trizol (Invitrogen, USA) and all total Nucleic Acid Isolation Kit (Ambion Inc., USA), following the manufacturer’s instruction. The experiment was performed at least three times using SYBR Premix Ex Taq (RR420A, TaKaRa, Japan) to quantify the mean values of delta Ct and SEM (Standard Error of Mean). The primers used for quantification were listed in Supplementary Table 1.

**Table S1:**
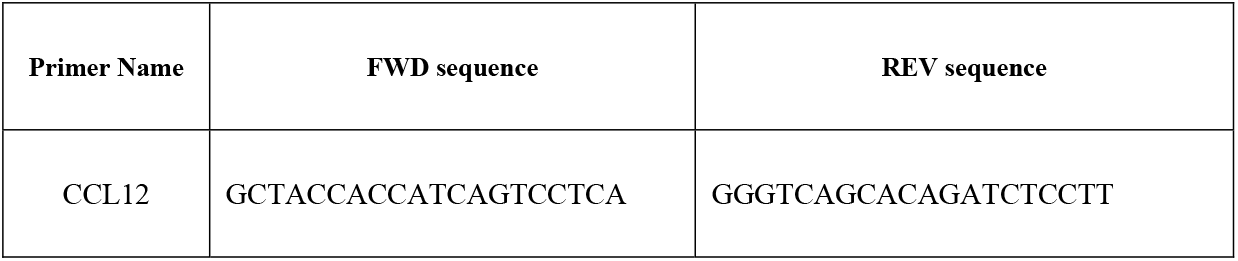
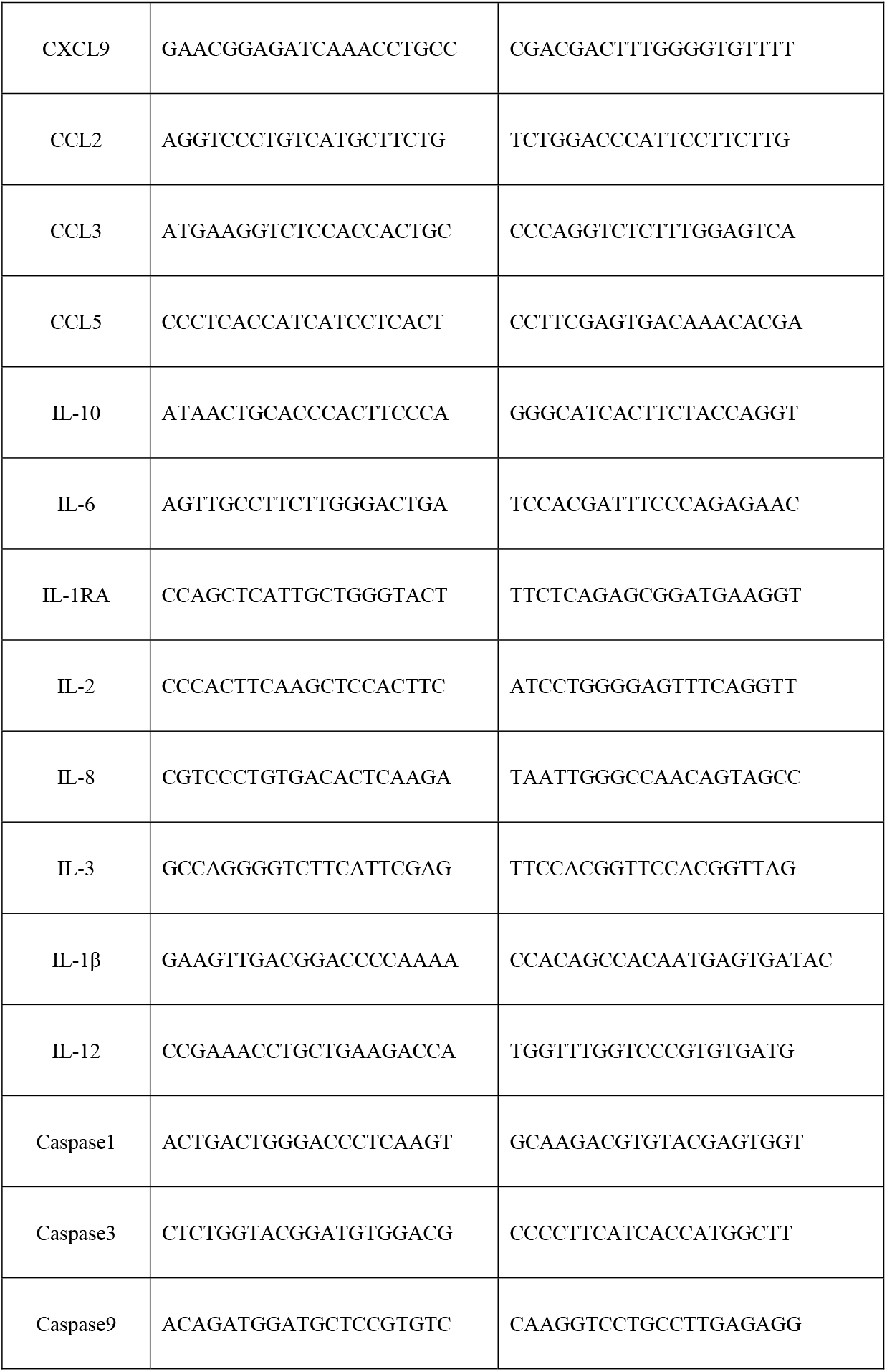
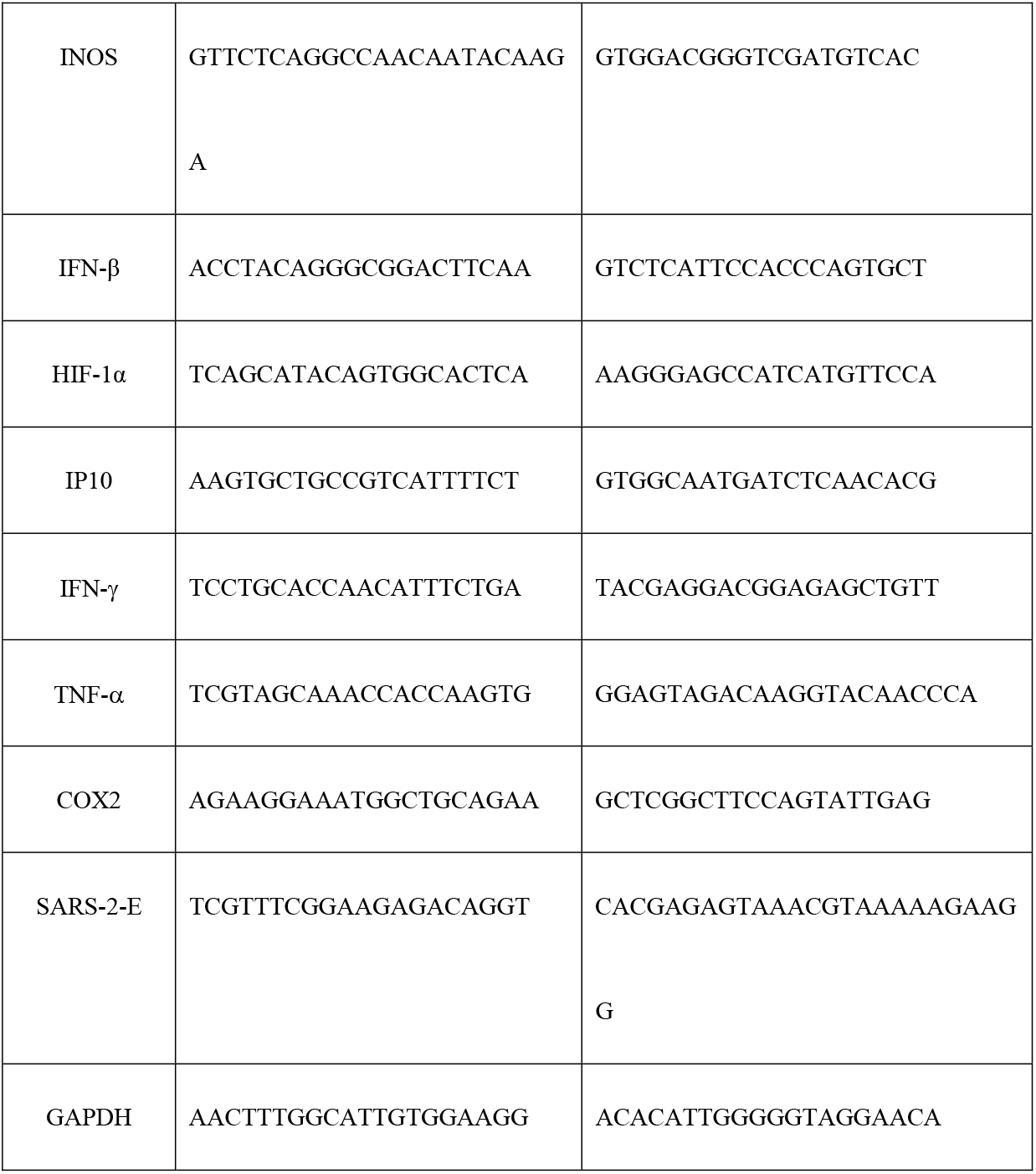
List of Quantitative Real-time PCR (qRT-PCR) primers.

### Protein purification

For expression of full length SARS-CoV-2-E protein, a 6× his tag was introduced at the C-terminus of SARS-CoV-2-E protein to facilitate purification. The pET28a vector containing recombinant protein was expressed in *E. coli* BL21 (DE3) pLysS competent cells (TransGen Biotech, China) and purified using Ni-NTA affinity chromatography (Yishen, China). Superdex 75 Increase 10/300 gel filtration chromatography (GE Healthcare, USA) was performed as the manufacturer’s instructions before concentration and application in animal model and electrophysiology recording.

### LC-MS/MS analysis

All LC-MS/MS experiments were carried on an online liquid chromatography-tandem mass spectrometry (LC-MS/MS) setup consisting of an Easy-nano LC system and a Q-Exactive HF mass spectrometer (Thermo, Bremen, Germany). The peptides were loaded on in-house packed column (75 μm × 200 mm fused silica column with 3 μm ReproSil-Pur C18 beads, Dr. Maisch GmbH, Ammerbuch, Germany) and separated with a 60-min gradient at a flow rate of 300 nL/min. Solvent A contained 100% H2O and 0.1% formic acid; Solvent B contained 100% acetonitrile and 0.1% formic acid. The spray voltage was set at 2,500 V in positive ion mode and the ion transfer tube temperature was set at 275°C. The MS1 full scan was set at a resolution of 60,000 at m/z 200, AGC target 3e6 and maximum IT 20 ms, followed by MS2 scans at a resolution of 15,000 at m/z 200, AGC target 1e5 and maximum IT 100 ms. The precursor ions were fragmented by higher energy collisional dissociation (HCD). Isolation window was set at 1.6 m/z. The normalized collision energy (NCE) was set at NCE 27%, and the dynamic exclusion time was 30 s. Precursors with charge 1, 7, 8 and > 8 were excluded for MS2 analysis.

The MS raw data were analyzed with MaxQuant (http://maxquant.org/, 1.6.7.0). Oxidized methionine and protein N-term acetylation were set as variable modifications. Trypsin/P or chymotrypsin was selected as the digestive enzyme with two potential missed cleavages. The false discovery rate (FDR) for peptides and proteins was rigorously controlled <1%.

### Animal model

SARS-Cov-2-E protein induced mouse model was established by conducting tail vein injection of purified protein (25 μg/g body weight) in 8-week-old mice. Male C57BL/6 mice were obtained from Shanghai SLAC Laboratory Animal Co., Ltd. All animal procedures were performed in accordance with the National Institutes of Health Guide for the Care and Use of Laboratory Animals, under protocols approved and strictly followed by the Institutional Animal Care and Use Committees (IACUC).

### Planar lipid bilayers recording

The purified proteins were incorporated into lipid bilayers to test their functionality. All the lipids were bought from Avanti. (Avanti Polar Lipids, USA). Proteins were added to *Cis* side and the buffer contained 500 mM KCl in *Cis/* 50 mM KCl in *Trans*, and all solutions were buffered by 5 mM HEPES, pH 6.35. Membrane currents were recorded under voltage-clamp mode using a Warnner bilayer clamp amplifier BC-535 (Warner Instruments, USA), filtered at 1-2 kHz. The currents were digitized using pClamp 10.2 software (Molecular Devices, US). Data are presented as means ± SEM. The single-channel conductance and open time were determined by fitting to Gaussian functions or to single or bi-exponential equations. Opening times less than 0.5 −1.5 ms were ignored. The equilibrium potential was calculated using the Nernst equation and Goldman-Hodgkin-Katz flux equation as reported (*33*).

### Viral inhibition assay

SARS-CoV-2 (nCoV-2019BetaCoV/Wuhan/WIV04/2019) was preserved at Wuhan institute of virology, Chinese Academy of Sciences. It was propagated and titrated with Vero E6 cells, and its associated operations were performed in a biosafety level 3 (BSL-3) facility. For viral inhibition assay, Vero E6 cells seeded in 48-well plate with 50000 cells per well overnight were firstly pre-treated with gradient-diluted compounds for 1 h, followed by adding SARS-CoV-2 (MOI = 0.01) and incubating for 1 h at 37°C. After that, the virus-drug mixture was completely removed, and the cells were washed twice with PBS followed by adding fresh medium with compounds. 24 h later, viral RNA was extracted from cell supernatants with Mini BEST Viral RNA/DNA Extraction Kit (Takara, Japan) according to the instructions, then reversely transcribed with Prime ScriptTM RT reagent Kit with gDNA Eraser (Takara, Japan). Viral genome copies were quantified with Takara TB Green^®^ Premix Ex Taq™ II (Takara, Japan) by a standard curve method on ABI 7500 using a pair of primers targeting S gene. The forward primer (5’-3’): GCTCCCTCTCATCAGTTCCA; the reverse primer (5’-3’): CTCAAGTGTCTGTGGATCACG.

### Inhibitor synthesis

#### General procedure

Reagents and solvents were purchased from commercial sources and used without purification. HSGF 254 (0.15–0.2 mm thickness) was used for analytical thin-layer chromatography (TLC). All products were characterized by their NMR and MS spectra. ^1^H NMR spectra was recorded on a Bruker spectrometer with TMS as the internal standard. Chemical shifts were given in δ values (ppm) and coupling constants (*J*) were given in Hz. Signal multiplicities were characterized as s (singlet), d (doublet), t (triplet), m (multiplet) and br (broad). The mass spectra were determined on a Thermo-Fisher Finnigan LTQ mass spectrometer. HPLC spectra were recorded by an Agilent 1100 (Agilent Corporation) with SunFire C18 (150 × 4.6 mm, 5 μm) with two solvent systems (acetonitrile/0.02 M 1-octanesulfonic acid sodium in water). Purity was determined by reversed-phase HPLC and was ≥ 95% for **BE30, BE31, BE32** and **BE33**.

**Scheme S1:**
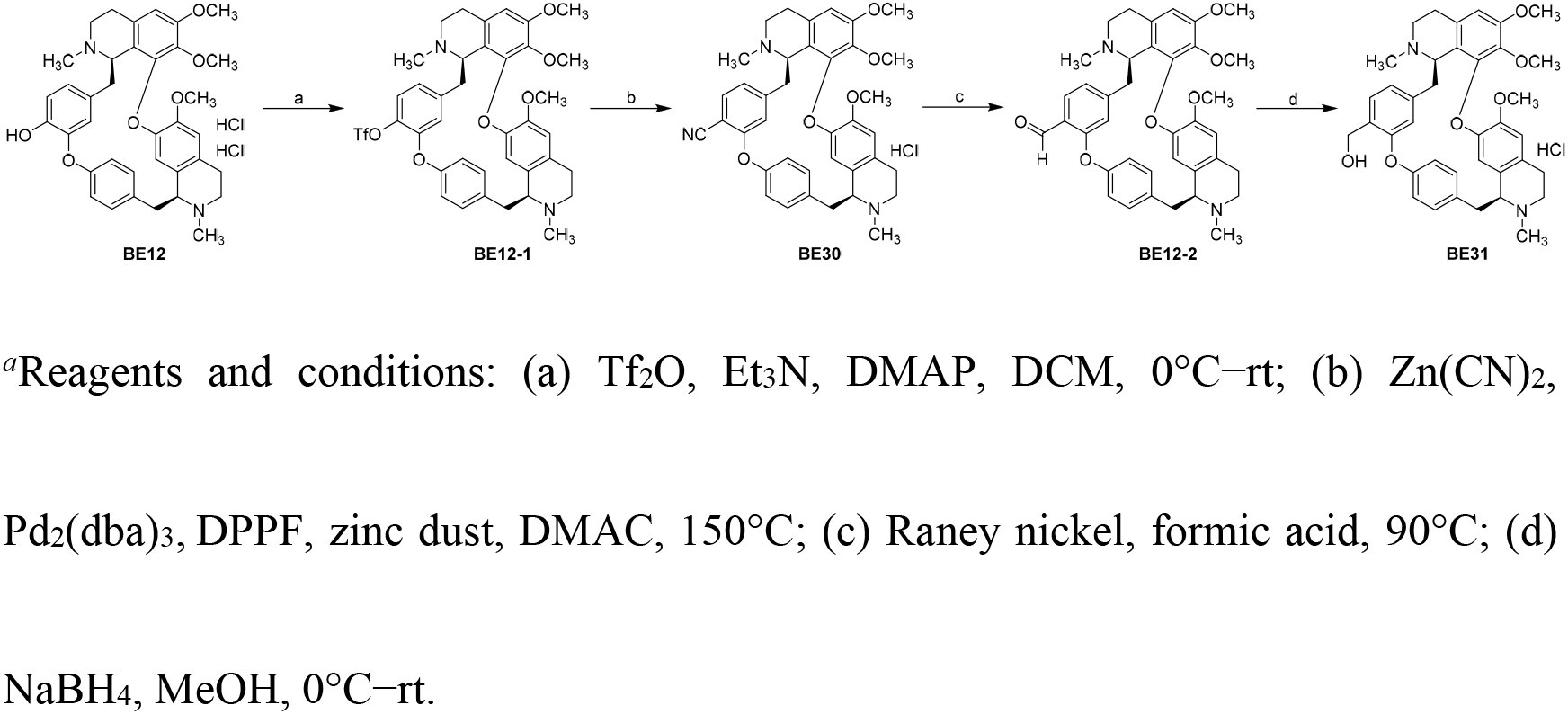
Synthetic route of compounds BE30 and BE31^*a*^

### Synthesis of BE12-1

To a 10 mL round-bottom flask was added phenol **BE12** (185 mg, 0.271 mmol), DCM (2 mL), DMAP (16.6 mg, 0.136 mmol, 0.5 eq) and triethylamine (0.189 mL, 1.355 mmol, 5 eq). The clear solution was stirred and cooled in an ice bath for 10 min. Triflic anhydride (0.091 mL, 0.542 mmol, 2 eq) was added dropwise over 2 mins. The reaction was slowly warmed to room temperature. After 30 mins, the reaction was diluted with DCM (5 mL) and quenched with 0.5 M aqueous HCl (1.1 mL). The organic layer was separated and washed with sat. aq. NaHCO_3_ (5 mL), then brine (5 mL), dried over Na_2_SO_4_, filtered, and concentrated via rotary evaporation. The crude product was purified by silica gel chromatography (DCM:MeOH = 80:1) to give triflate **BE12-1** as a light-yellow solid (160 mg, 0.217 mmol, 80% yield). ^1^H NMR (500 MHz, Chloroform-*d*) δ 7.47 (dd, *J* = 8.5, 2.2 Hz, 1H), 7.07 (d, *J* = 8.3 Hz, 1H), 6.97 (td, *J* = 8.8, 2.4 Hz, 2H), 6.83 (dd, *J* = 8.4, 2.0 Hz, 1H), 6.61 (s, 1H), 6.37 (s, 1H), 6.34-6.32 (m, 1H), 6.31 (s, 1H), 5.60 (d, *J* = 2.0 Hz, 1H), 4.20 (d, *J* = 5.9 Hz, 1H), 3.79 (s, 3H), 3.66 (d, *J* = 3.4 Hz, 1H), 3.63 (s, 3H), 3.38 (d, *J* = 14.4 Hz, 1H), 3.24-3.14 (m, 5H), 3.05 (m, 1H), 2.94 (m, 1H), 2.83 (m, 2H), 2.73 (m, 2H), 2.67 (s, 3H), 2.56 (s, 3H), 2.41 −2.29 (m, 3H). ESI-MS (*m/z*): 741.4 [M + H]^+^.

### Synthesis of BE30

A mixture of triflate **BE12-1** (170 mg, 0.230 mmol), Zn(CN)_2_ (40 mg, 0.345 mmol, 1.5 eq), Pd_2_(dba)_3_ (11 mg, 0.012 mmol, 0.05 eq), 1,1’-bis(diphenylphosphino)ferrocene (13 mg, 0.023 mmol, 0.1 eq) and zinc dust (2 mg, 0.03 mmol, 0.13 eq) in DMAC (2 mL) in a 10 mL round-bottom flask was purged with nitrogen for three times at room temperature. Then the mixture was heated to 150 °C and stirred at this temperature for 3 h. After cooling to room temperature, ethyl acetate (10 mL) was added and the resulting mixture was filtered. The filtrate was washed with water (2 × 5 mL), brine (5 mL), dried over Na_2_SO_4_, filtered, and concentrated via rotary evaporation. The crude product was purified by silica gel chromatography (DCM:MeOH = 80:1) to give free base of **BE30** as a light-yellow solid (110 mg, 0.178 mmol, 78% yield). ^1^H NMR (500 MHz, Chloroform-*d*) δ 7.49 (dd, *J* = 8.4, 2.3 Hz, 1H), 7.39 (d, *J* = 7.9 Hz, 1H), 6.99 (dd, *J* = 8.2, 2.3 Hz, 1H), 6.95 (dd, *J* = 8.3, 2.6 Hz, 1H), 6.87 (dd, *J* = 7.9, 1.4 Hz, 1H), 6.60 (s, 1H), 6.37 (s, 1H), 6.33 (dd, *J* = 8.3, 2.6 Hz, 1H), 6.31 (s, 1H), 5.55 (d, *J* = 1.4 Hz, 1H), 4.22 (d, *J* = 6.0 Hz, 1H), 3.79 (s, 3H), 3.69 (d, *J* = 3.8 Hz, 1H), 3.64 (s, 3H), 3.44-3.39 (m, 1H), 3.28-3.14 (m, 5H), 3.06 (m, 1H), 2.96 (m, 1H), 2.88-2.79 (m, 2H), 2.78-2.70 (m, 2H), 2.68 (s, 3H), 2.56 (s, 3H), 2.40-2.27 (m, 3H). To a solution of free base of **BE30** (105 mg) in methanol (1 mL) was added concentrated hydrochloric acid (14.9 μL). The resulting solution was stirred at room temperature for 5 min and then concentrated via rotary evaporation. Methyl tert-butyl ether (1 mL) was added to the residue and the precipitation was filtered and dried under vacuum at 45 °C to provide hydrochloride salt of **BE30** as a yellow solid (90 mg). ESI-MS (*m/z*): 618.4 [M + H]^+^.

### Synthesis of BE12-2

A mixture of hydrochloride salt of **BE30** (75 mg, 0.115 mmol) and Raney nickel (100 mg) was suspended in formic acid (3 mL) and water (0.1 mL) and heated to 90 °C for 2 h. The mixture was then filtered, rinsed with methanol, and concentrated. To the resulting residue was added DCM (5 mL) followed by sat. aq. NaHCO_3_ until basic. The organic layer was separated, dried over Na_2_SO_4_, and concentrated. The crude product was purified by silica gel chromatography (DCM:MeOH = 75:1) to give aldehyde **BE12-2** as a light-yellow oil (38 mg, 0.061 mmol, 57% yield). ESI-MS (*m/z*): 621.4 [M + H]^+^.

### Synthesis of BE31

Sodium borohydride (4 mg, 0.106 mmol, 1.8 eq) was added to a solution of **BE12-2** (36 mg, 0.058 mmol) in methanol (1 mL) under an ice bath, and the solution was slowly warmed to room temperature. After 2 h, the reaction was quenched with sat. aq. NH_4_Cl (1 mL) and extracted with DCM (2 × 2 mL). The combined organic layer was washed with brine (2 mL), dried over Na_2_SO_4_, filtered, and concentrated via rotary evaporation. The crude product was purified by silica gel chromatography (DCM:MeOH = 65:1) to give free base of **BE31** as a light-yellow solid (20 mg, 0.032 mmol, 55% yield). Two sets of ^1^H NMR data representing two rotamers (~3:2) were observed as indicative of the presence of atropisomers in bisbenzylisoquinoline type alkaloids involving planar chirality. ^1^H NMR (600 MHz, Chloroform-*d*, major rotamer) δ 7.80 (brs, 1H), 7.37 (d, *J* = 8.4 Hz, 1H), 7.21 (d, *J* = 7.3 Hz, 1H), 6.92 (m, 2H), 6.68 (s, 1H), 6.44 (s, 1H), 6.37 (s, 1H), 6.32 (dd, *J* = 8.1, 2.6 Hz, 1H), 5.32 (s, 1H), 4.78 (m, 2H), 4.59 (m, 1H), 4.14–3.85 (m, 2H), 3.82 (s, 3H), 3.78–3.46 (m, 2H), 3.65 (s, 3H), 3.36–3.13 (m, 3H), 3.22 (s, 3H), 3.08–2.91 (m, 3H), 2.90–2.67 (m, 3H), 2.82 (s, 3H), 2.73 (s, 3H); ^1^H NMR (600 MHz, Chloroform-*d*, minor rotamer) δ 7.47 (d, *J* = 7.8 Hz, 1H), 7.43 (brs, 1H), 7.16 (brs, 1H), 6.92 (m, 2H), 6.61 (dd, *J* = 8.5, 2.7 Hz, 1H), 6.48 (s, 1H), 6.47 (s, 1H), 6.44 (s, 1H), 5.32 (s, 1H), 4.78 (m, 2H), 4.75 (m, 1H), 4.14–3.85 (m, 2H), 3.83 (s, 3H), 3.78–3.46 (m, 2H), 3.43 (s, 3H), 3.36–3.13 (m, 3H), 3.18 (s, 3H), 3.08–2.91 (m, 3H), 2.90–2.67 (m, 3H), 2.79 (s, 3H), 2.69 (s, 3H). To a solution of free base of **BE31** (16 mg) in methanol (0.5 mL) was added concentrated hydrochloric acid (2.2 μL). The resulting solution was stirred at room temperature for 5 mins and then concentrated via rotary evaporation. Acetonitrile (0.5 mL) was added to the residue and the precipitation was filtered and dried under vacuum at 45 °C to provide hydrochloride salt of **BE31** as an off-white solid (13 mg). ESI-MS (*m/z*): 623.4 [M + H]^+^.

**Scheme S2:**
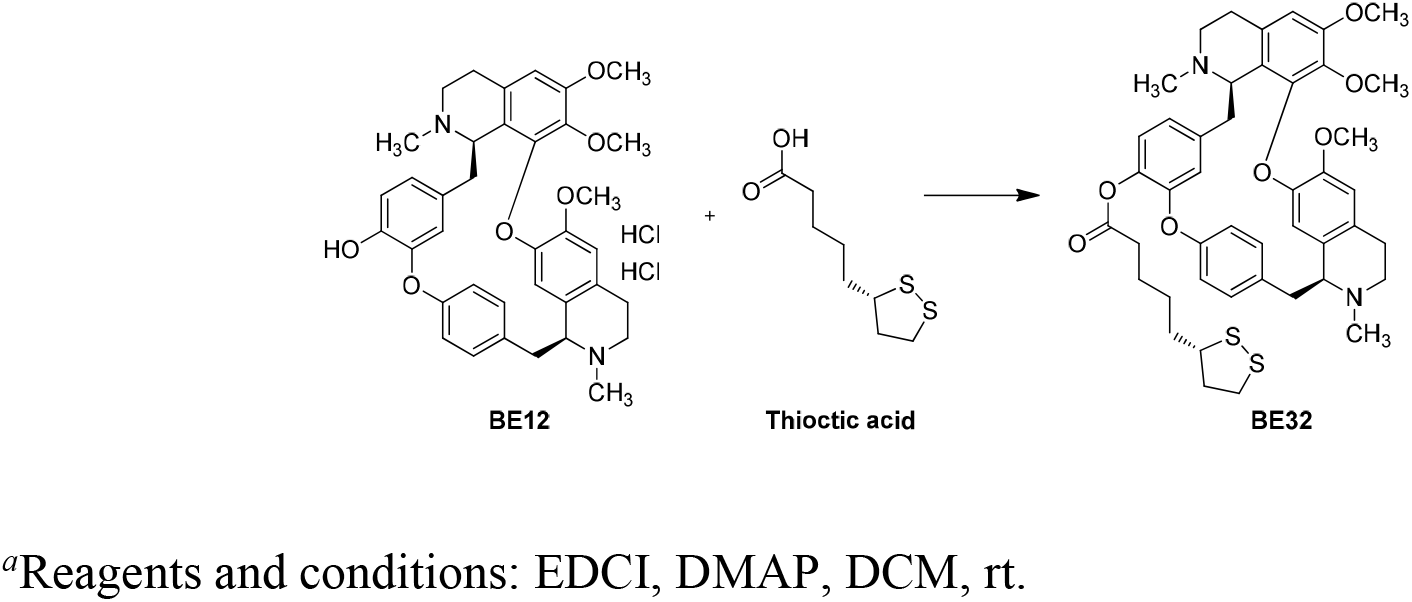
Synthetic route of compound BE32^*a*^

### Synthesis of BE32

EDCI (43 mg, 0.224 mmol, 1.5 eq) and DMAP (4 mg, 0.033 mmol, 0.2 eq) were added to a suspension of phenol **BE12** (100 mg, 0.147 mmol) and L-thioctic acid (31 mg, 0.150 mmol, 1.0 eq) in DCM (3 mL) at room temperature. The mixture was stirred for 5 h before water (2 mL) was added. The organic layer was separated, washed with brine (2 mL), dried over Na_2_SO_4_, filtered, and concentrated. The crude product was purified by silica gel chromatography (DCM:MeOH = 50:1) to give **BE32** as a light-yellow solid (63 mg, 0.079 mmol, 54% yield). Two sets of ^1^H NMR data representing two rotamers (~7:3) were observed. ^1^H NMR (500 MHz, Methanol-*d_4_*, major rotamer) δ 7.45 (dd, *J* = 8.5, 2.2 Hz, 1H), 7.16 (m, 1H), 6.99 (dd, *J* = 8.3, 2.2 Hz, 1H), 6.90 (m, 2H), 6.74 (s, 1H), 6.51 (s, 1H × 2), 6.34 (dd, *J* = 8.2, 2.6 Hz, 1H), 5.59 (d, *J* = 1.7 Hz, 1H), 4.25 (d, *J* = 5.7 Hz, 1H), 3.80 (s, 3H), 3.75 (d, *J* = 3.7 Hz, 1H), 3.64 (s, 3H), 3.46 (m, 1H), 3.24–3.16 (m, 2H), 3.20 (s, 3H), 3.10–3.04 (m, 2H), 3.03–2.96 (m, 2H), 2.93 (m, 1H), 2.82–2.70 (m, 3H), 2.63–2.37 (m, 5H), 2.60 (s, 3H), 2.57 (s, 3H), 2.32 (m, 1H), 1.80–1.48 (m, 8H); ^1^H NMR (500 MHz, Methanol-*d_4_*, minor rotamer) δ 7.34 (dd, *J* = 8.5, 2.2 Hz, 1H), 7.16 (m, 1H), 6.90 (m, 2H), 6.79 (dd, *J* = 8.4, 2.7 Hz, 1H), 6.69 (s, 1H), 6.55 (s, 1H), 6.54 (s, 1H), 6.37 (m, 1H), 4.31 (dd, *J* = 11.4, 4.2 Hz, 1H), 3.80 (m, 4H), 3.46 (m, 2H), 3.40 (s, 3H), 3.24–3.16 (m, 2H), 3.13 (s, 3H), 3.10–3.04 (m, 2H), 3.03–2.96 (m, 2H), 2.93 (m, 1H), 2.82–2.70 (m, 3H), 2.63–2.37 (m, 5H), 2.59 (s, 3H), 2.52 (s, 3H), 2.32 (m, 1H), 1.80–1.48 (m, 8H). ESI-MS (*m/z*): 797.5 [M + H]^+^.

**Scheme S3:**
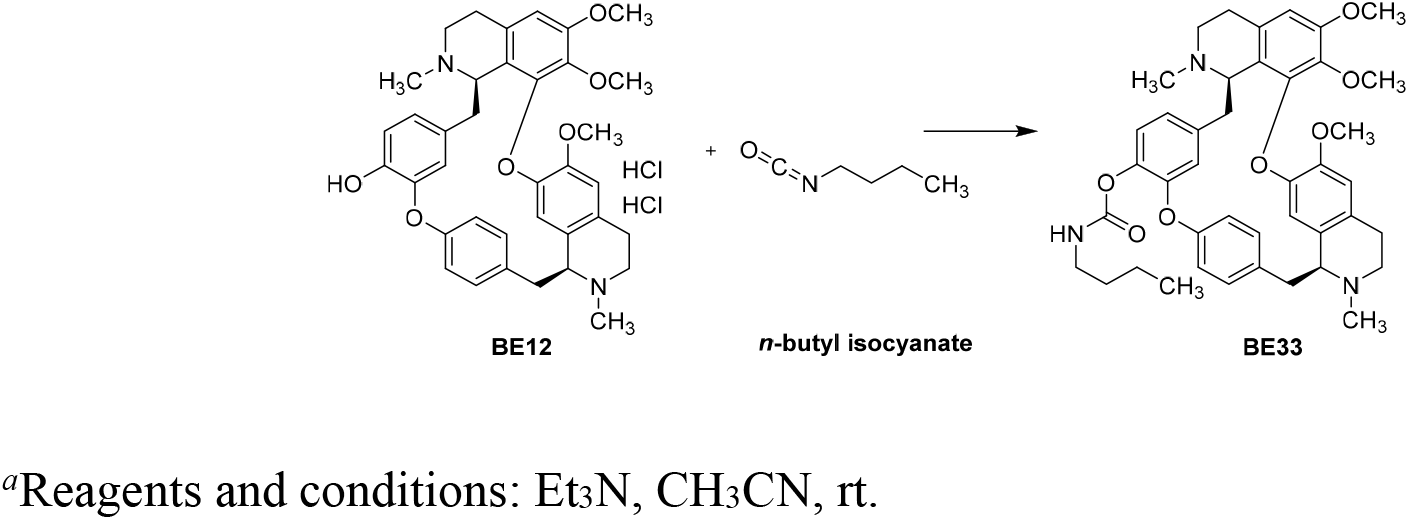
Synthetic route of compound BE33^*a*^

### Synthesis of BE33

Triethylamine (0.073 mL, 0.524 mmol, 3.6 eq) was added to a suspension of phenol **BE12** (100 mg, 0.147 mmol) and *n*-butyl isocyanate (22 mg, 0.222 mmol, 1.5 eq) in acetonitrile (3 mL) at room temperature. The mixture was stirred for 2 h, concentrated and subjected to silica gel chromatography (DCM:MeOH=30:1) to give **BE33** as a light-yellow solid (72 mg, 0.102 mmol, 69% yield). Two sets of ^1^H NMR data representing two rotamers (~7:3) were observed. ^1^H NMR (500 MHz, Methanol-*d_4_*, major rotamer) δ 7.45 (dd, *J* = 8.4, 2.3 Hz, 1H), 7.13 (m, 1H), 6.98 (dd, *J* = 7.9, 2.1 Hz, 1H), 6.94 (d, *J* = 8.1 Hz, 1H), 6.89 (m, 1H), 6.76 (s, 1H), 6.53 (s, 1H), 6.52 (s, 1H), 6.37 (dd, *J* = 8.2, 2.6 Hz, 1H), 5.53 (d, *J* = 1.9 Hz, 1H), 4.34 (d, *J* = 5.9 Hz, 1H), 3.92 (m, 1H), 3.80 (s, 3H), 3.64 (s, 3H), 3.37 (m, 1H), 3.24 (m, 1H), 3.19 (s, 3H), 3.17 (m, 3H), 3.06 (m, 2H), 2.92–2.76 (m, 4H), 2.72–2.42 (m, 3H), 2.67 (s, 3H), 2.65 (s, 3H), 1.49 (m, 2H), 1.33 (m, 2H), 0.89 (t, *J* = 7.3 Hz, 3H); ^1^H NMR (500 MHz, Methanol-*d_4_*, minor rotamer) δ 7.35 (dd, *J* = 8.5, 2.3 Hz, 1H), 7.19 (d, *J* = 8.2 Hz, 1H), 6.89 (m, 2H), 6.83 (dd, *J* = 8.4, 2.7 Hz, 1H), 6.69 (s, 1H), 6.59 (s, 1H), 6.56 (s, 1H), 6.39 (m, 1H), 4.50 (dd, *J* = 11.5, 4.2 Hz, 1H), 3.80 (m, 4H), 3.67 (m, 1H), 3.49 (d, *J* = 7.3 Hz, 1H), 3.40 (s, 3H), 3.24 (m, 1H), 3.17 (m, 3H), 3.10 (s, 3H), 3.06 (m, 2H), 2.92–2.76 (m, 4H), 2.72–2.42 (m, 3H), 2.66 (s, 3H), 2.61 (s, 3H), 1.49 (m, 2H), 1.33 (m, 2H), 0.86 (t, *J* = 7.3 Hz, 3H). ESI-MS (*m/z*): 708.5 [M + H]^+^.

### Supplemental Video and Figures

#### Video S1

16HBE cells were recorded 7 h after transfected with 2-E-GFP for the duration of time indicated on the left corner (h : min : s: ms). Scale bar, 25 μm. Also see Figure 1.

**Figure S1.**
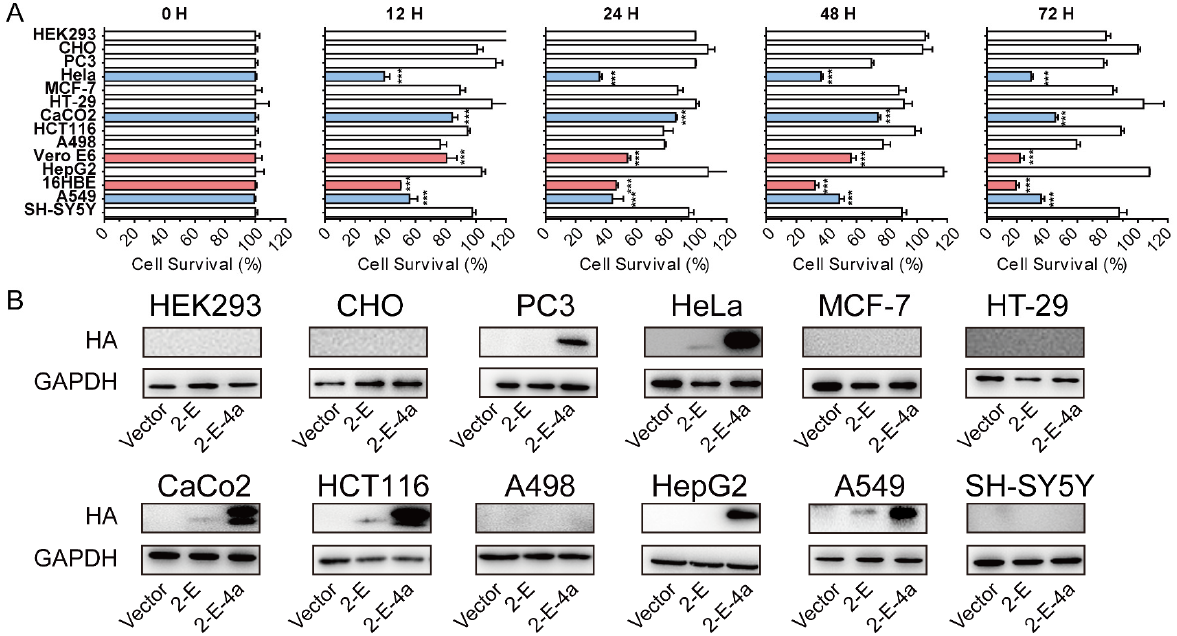
Cell viability and expression after 2-E transfection. A. Cell viability of 14 cell lines at indicated time after transfecting with 2-E-4a plasmids. B. Expression of 2-E and 2-E-4a in various cell lines.

**Figure S2.**
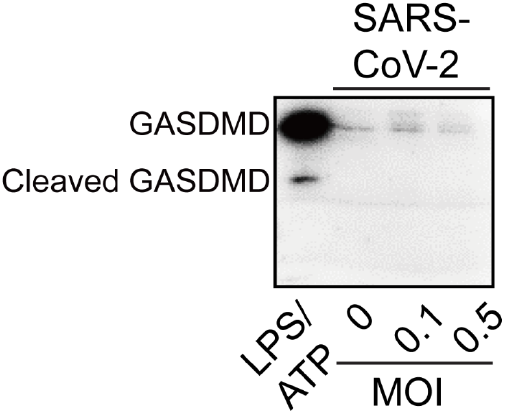
Western blot showed the expression of cleaved GSDMD after treatment of Lipopolysaccharide (LPS) 10 ng/mL plus Adenosine Triphosphate (ATP) 5 mM in macrophages (Wang et al., 2020).

**Figure S3.**
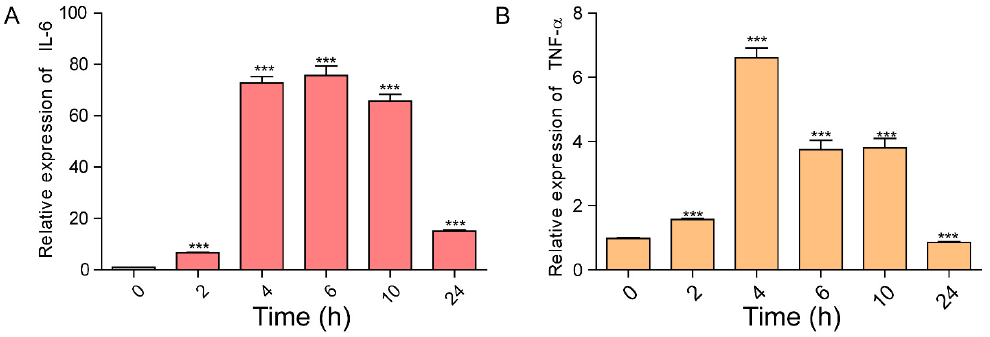
Expression levels of IL-6 and TNF-α at indicated time after transfection with 2-E plasmids.

**Figure S4.**
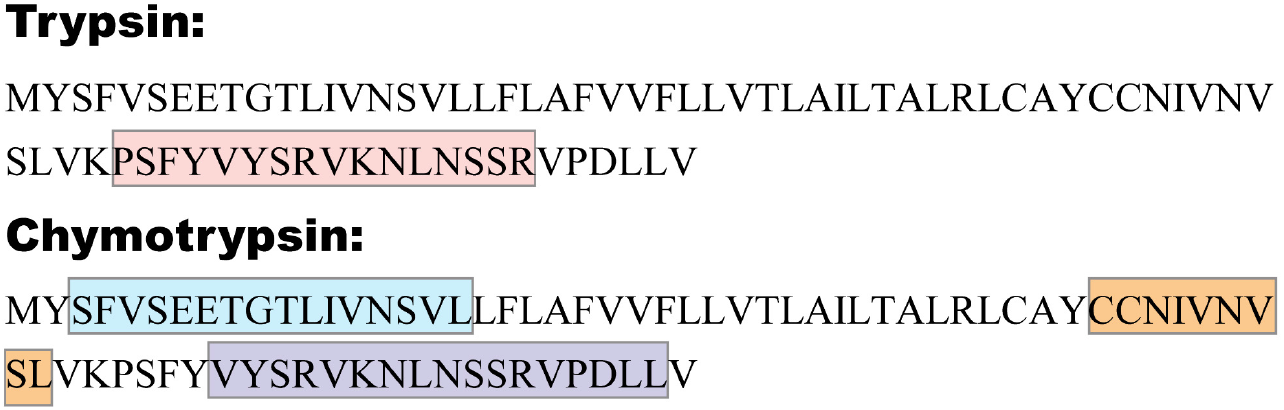
Four pieces of peptides (red, blue, orange and purple) were detected using LC-MS/MS when the purified 2-E proteins were treated with trypsin or chymotrypsin, respectively.

**Figure S5.**
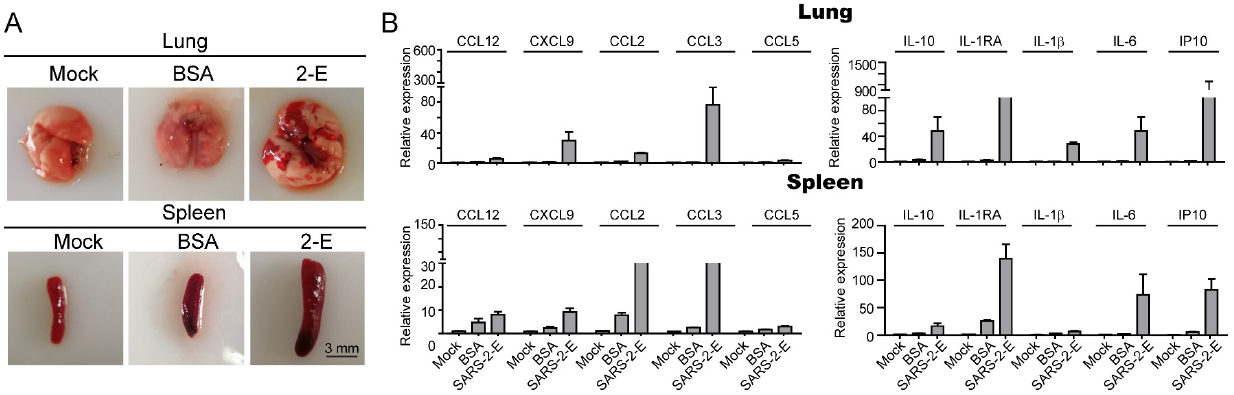
Bovine serum albumin did not cause acute respiratory distress syndrome (ARDS)-like damage in the lung and spleen in mice. A. Gross pathology and histopathology of lung and spleen from control mice (Trisbuffered saline, TBS; Bovine serum albumin, BSA, (25 μg/g body weight)), and model mice (2-E proteins). B. qRT-PCR analysis of the lung and spleen tissues after injection of TBS, BSA and 2-E proteins.

**Figure S6:**
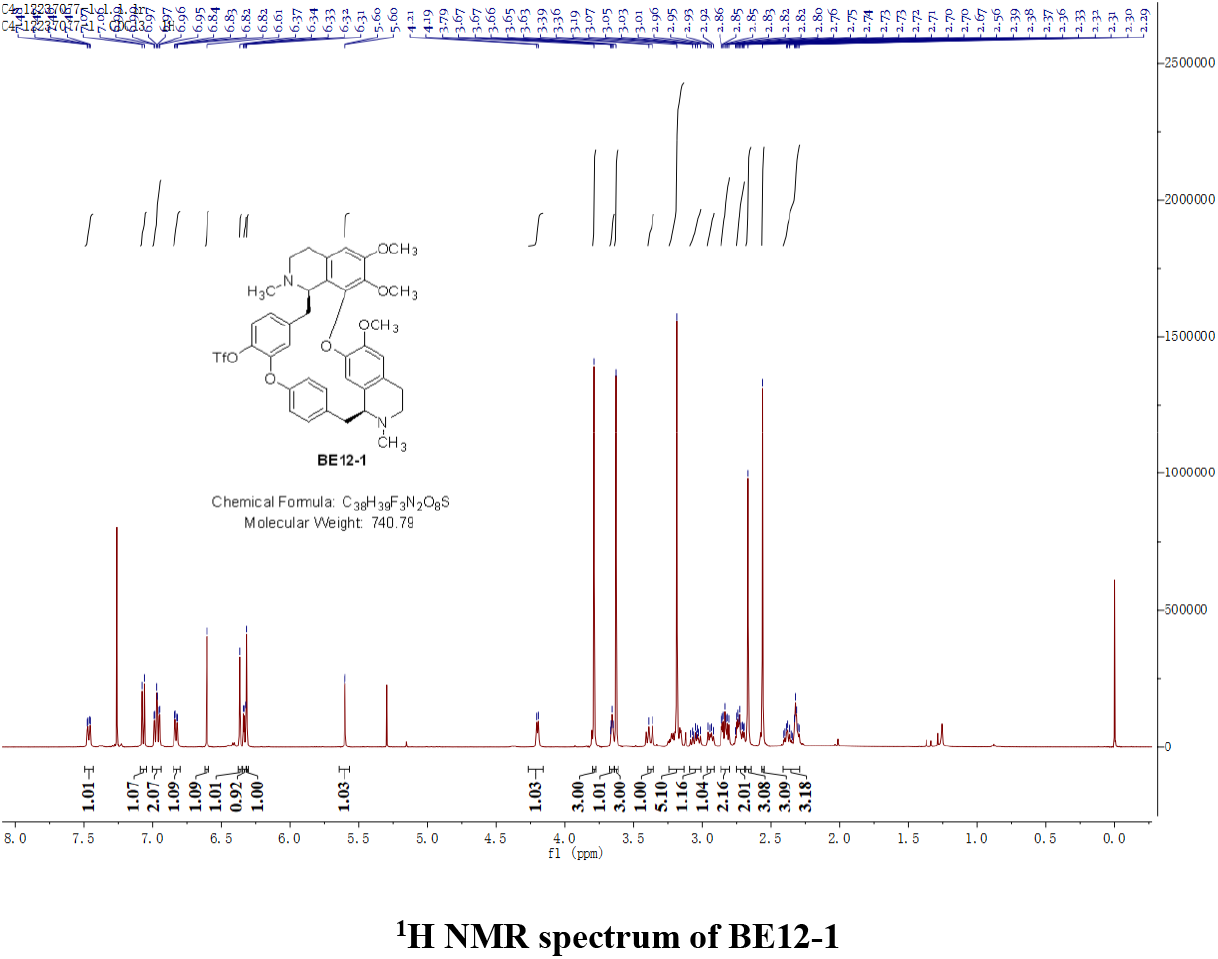
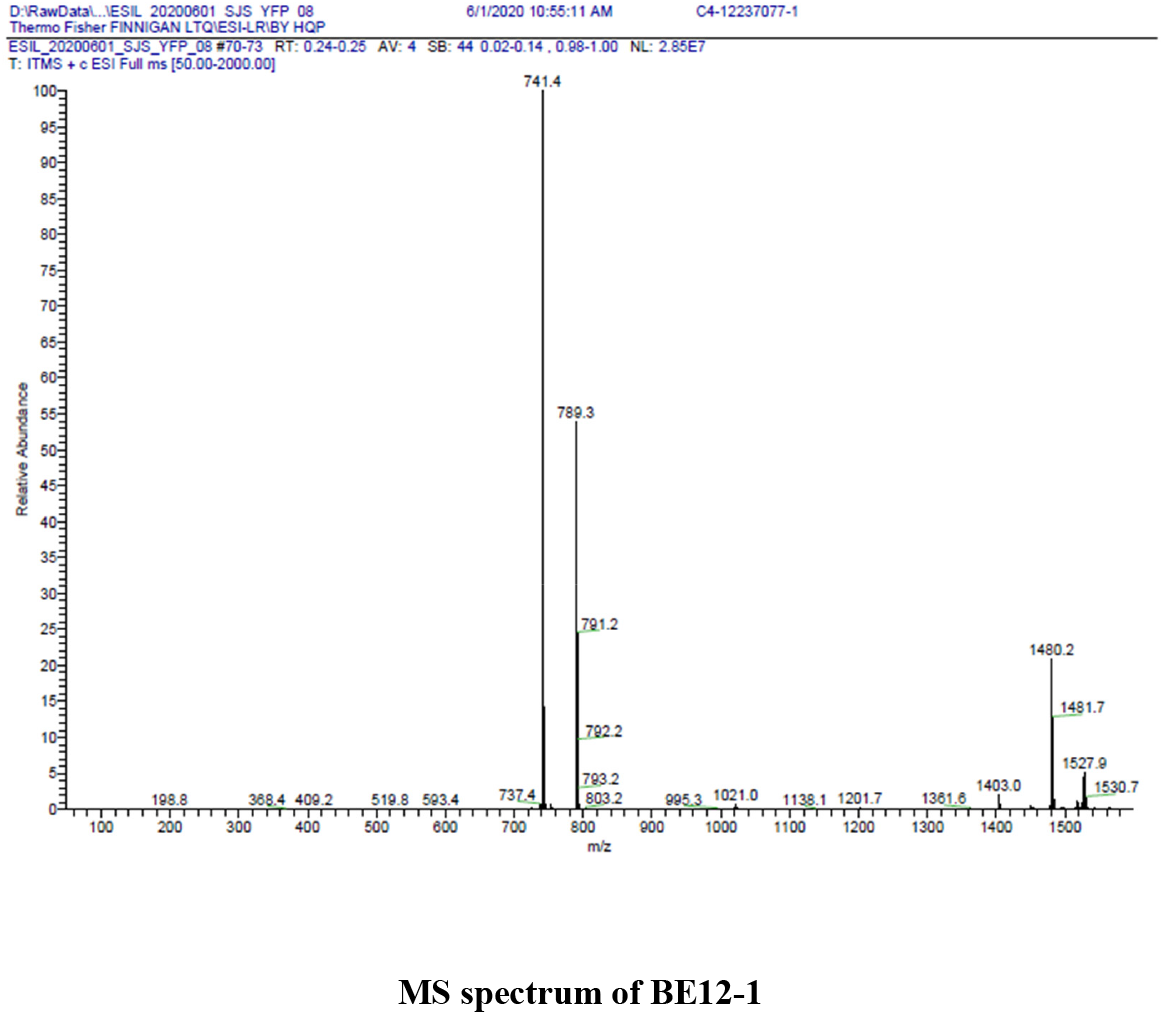
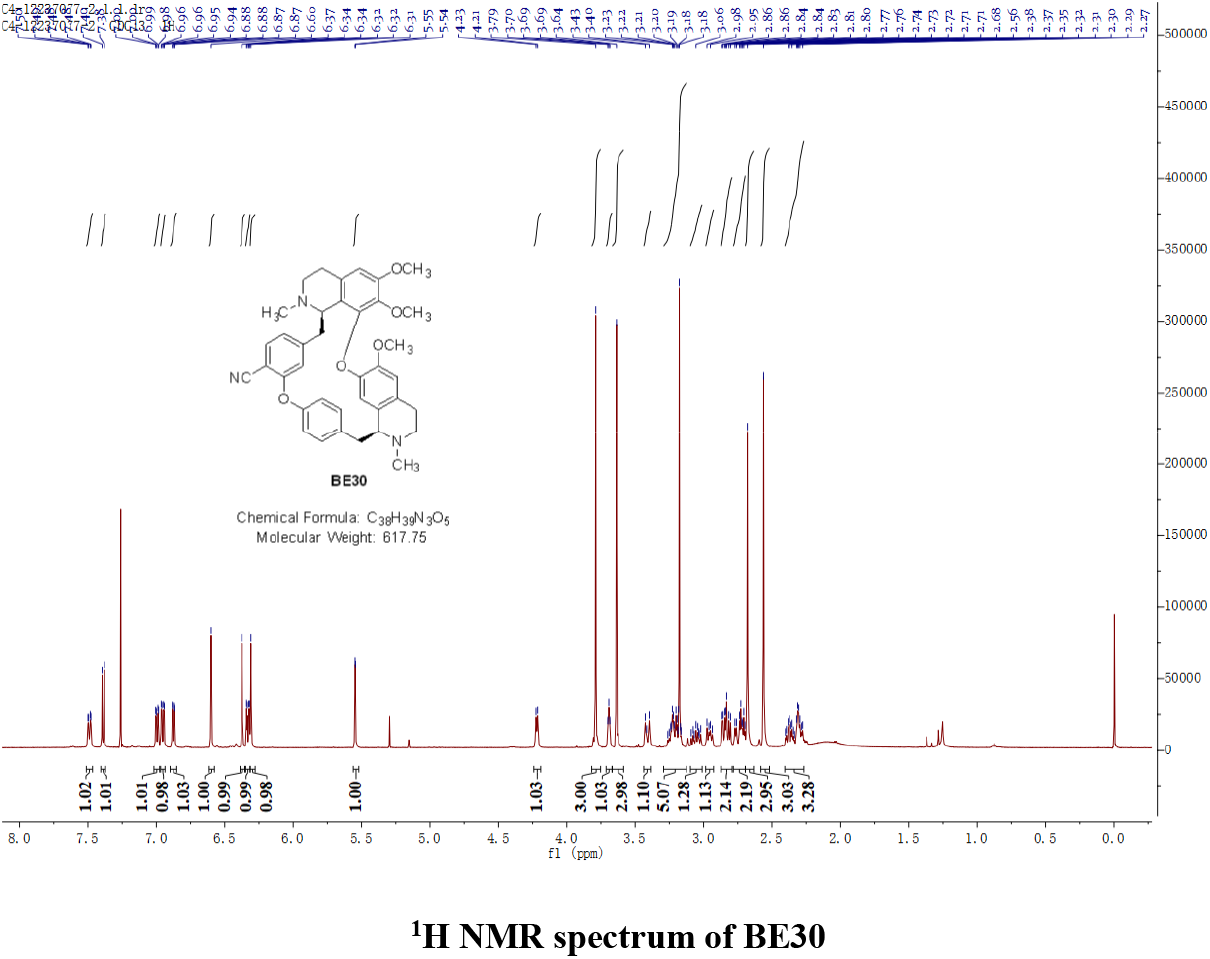
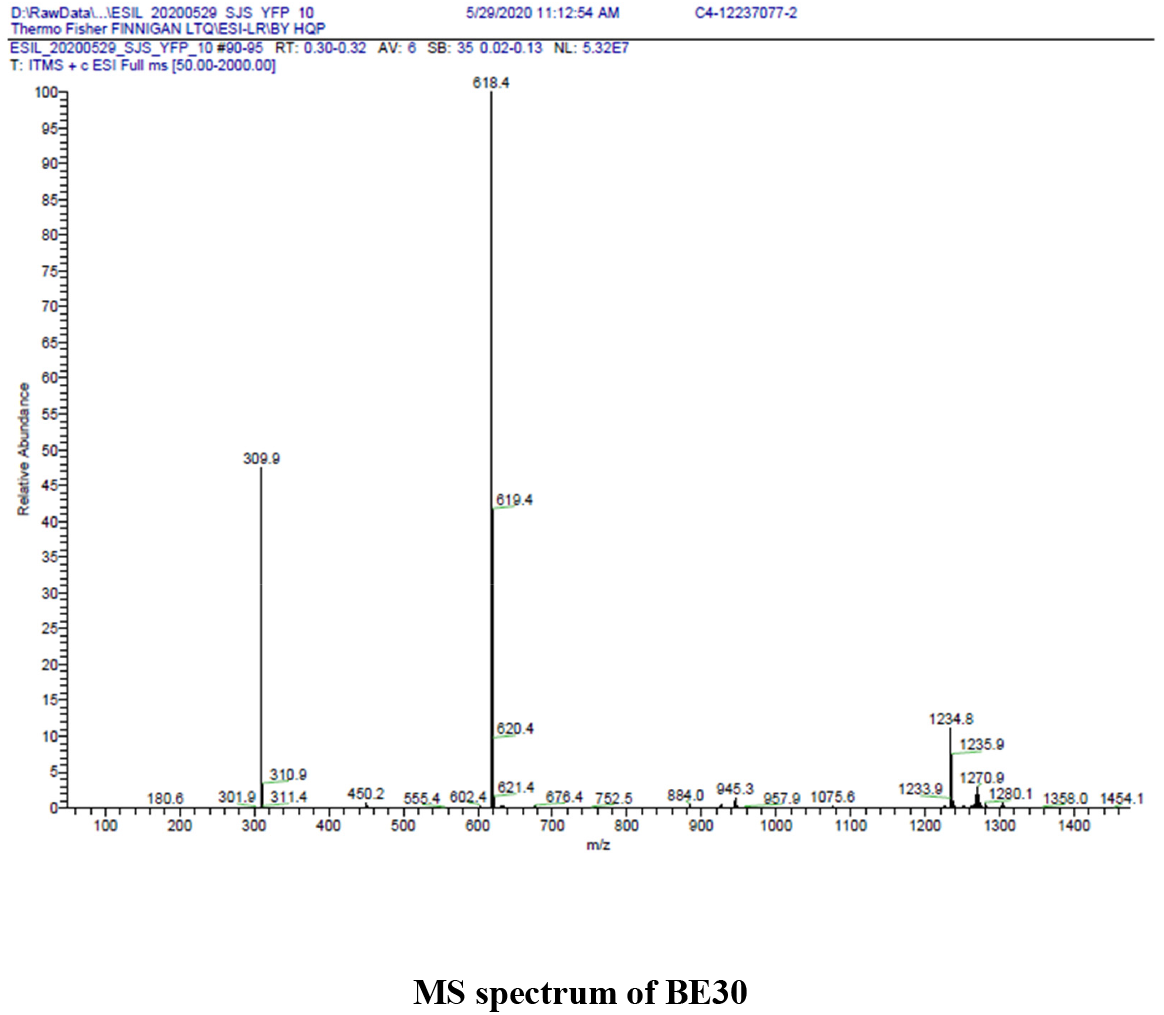
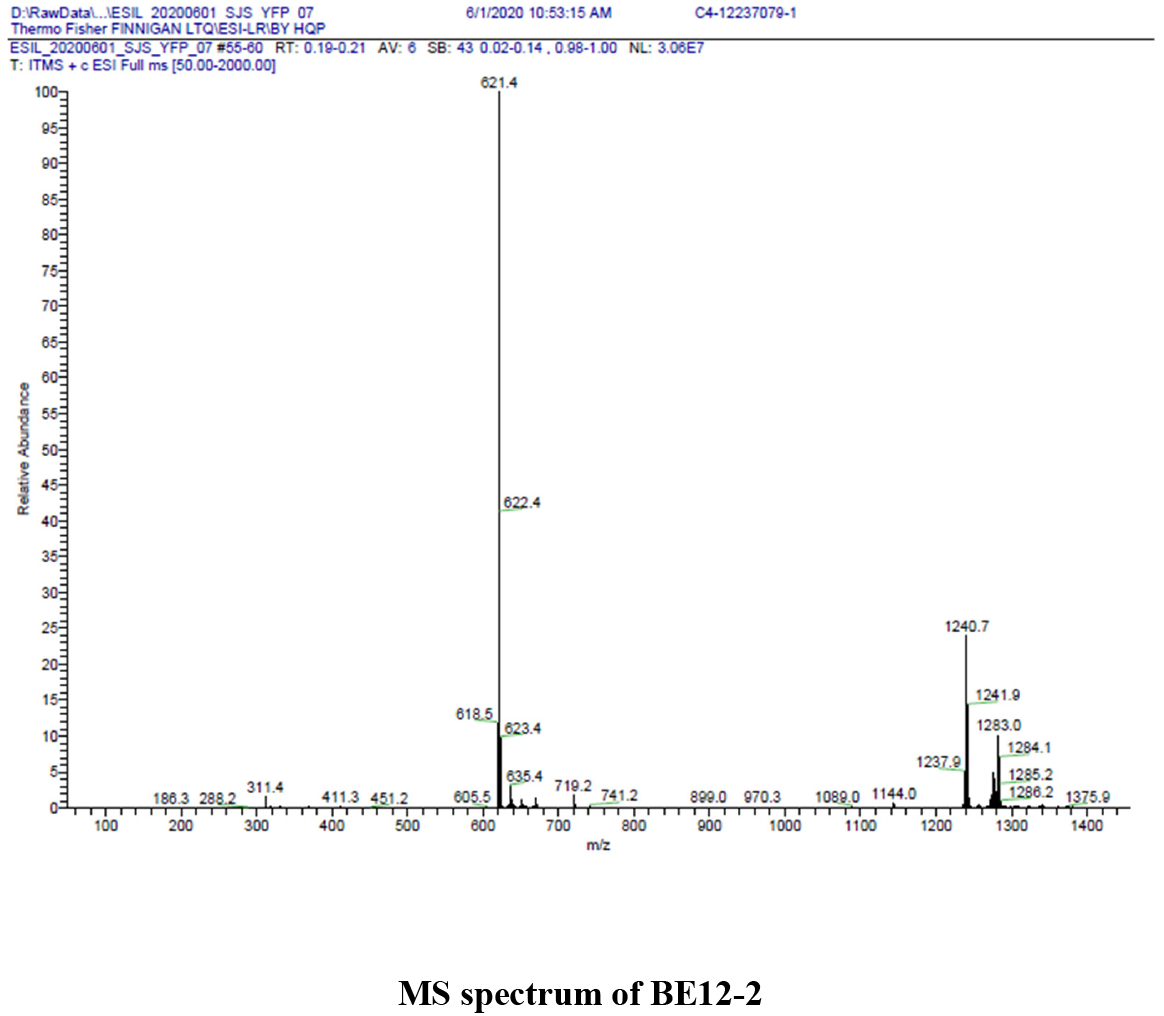
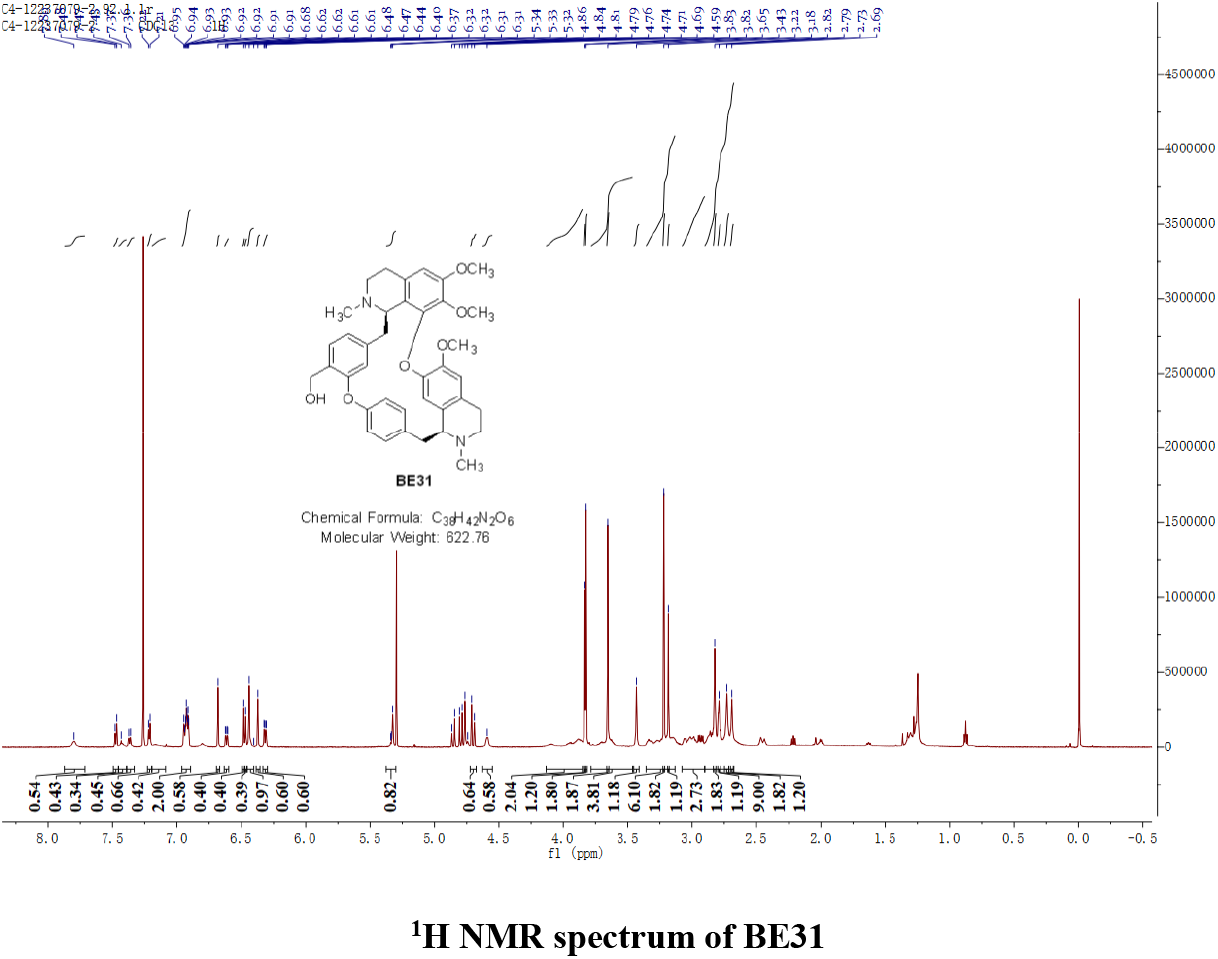
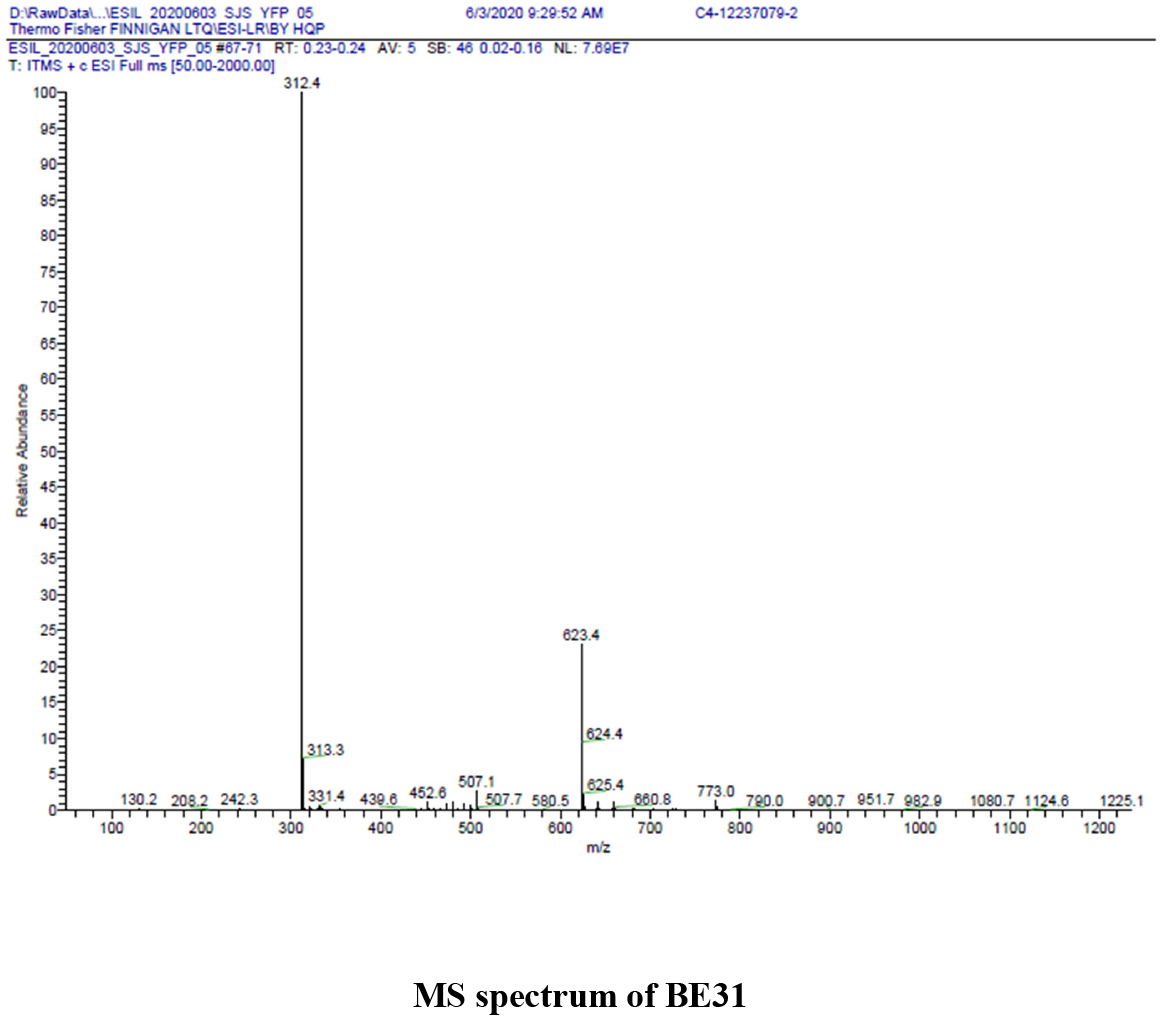
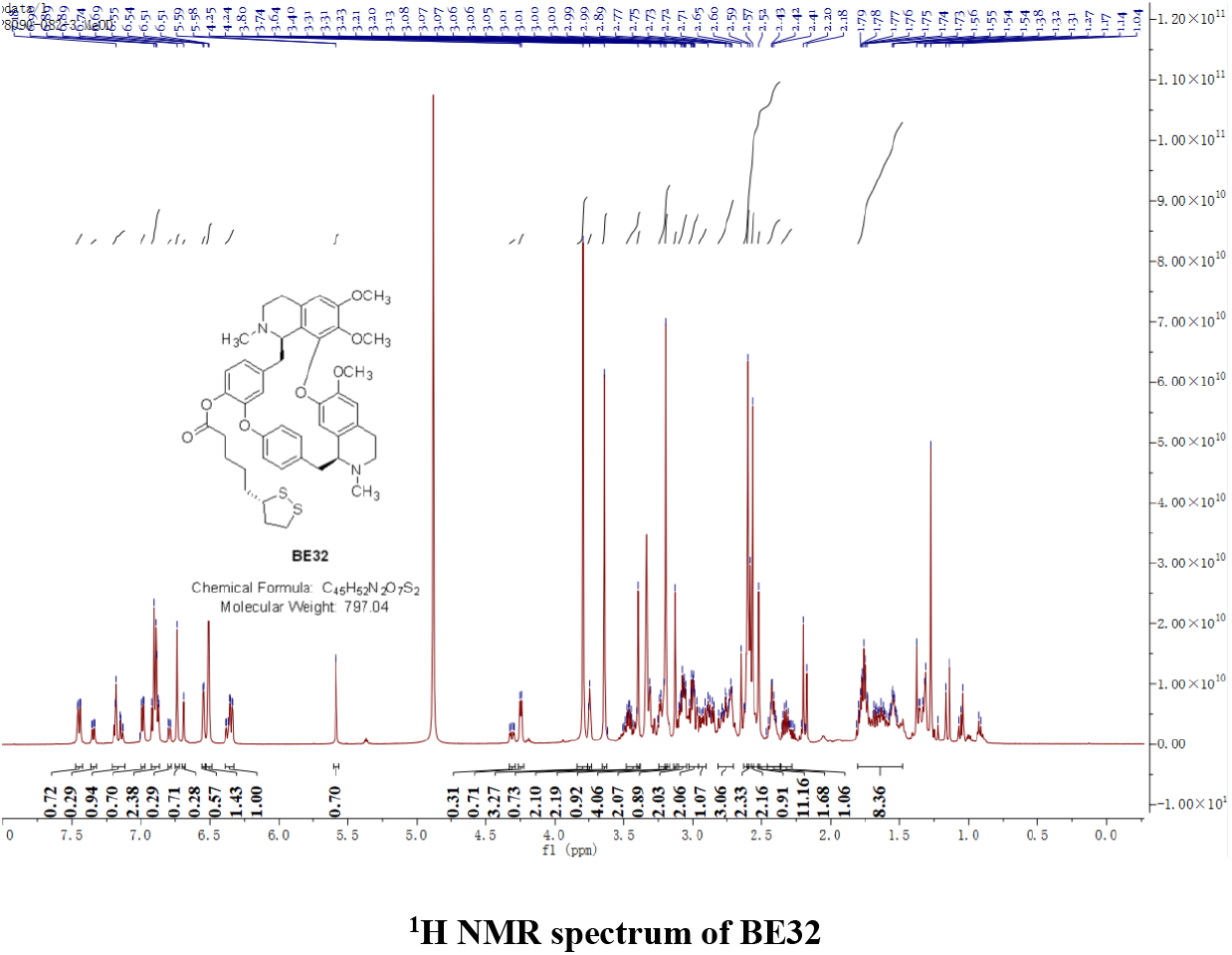
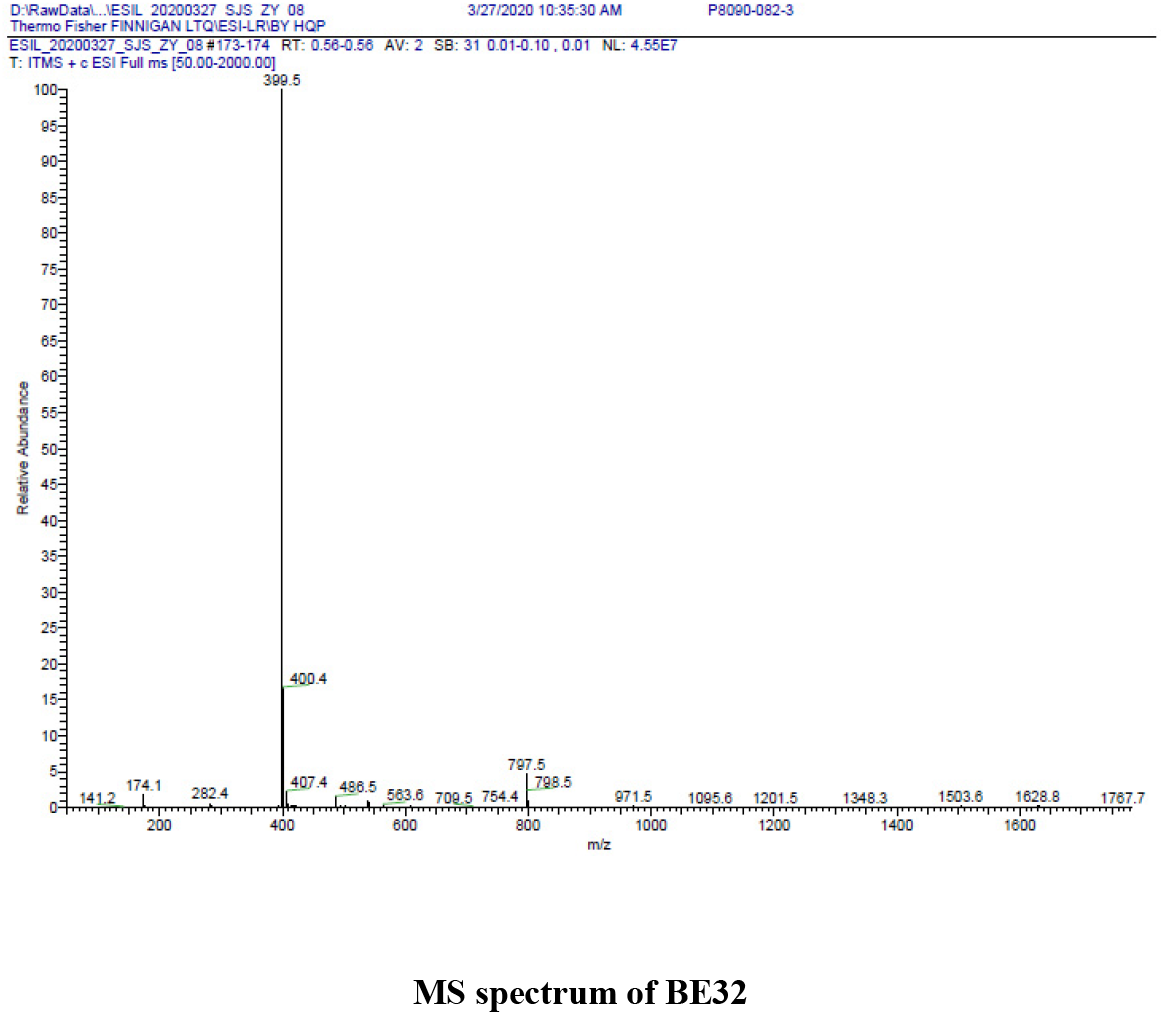
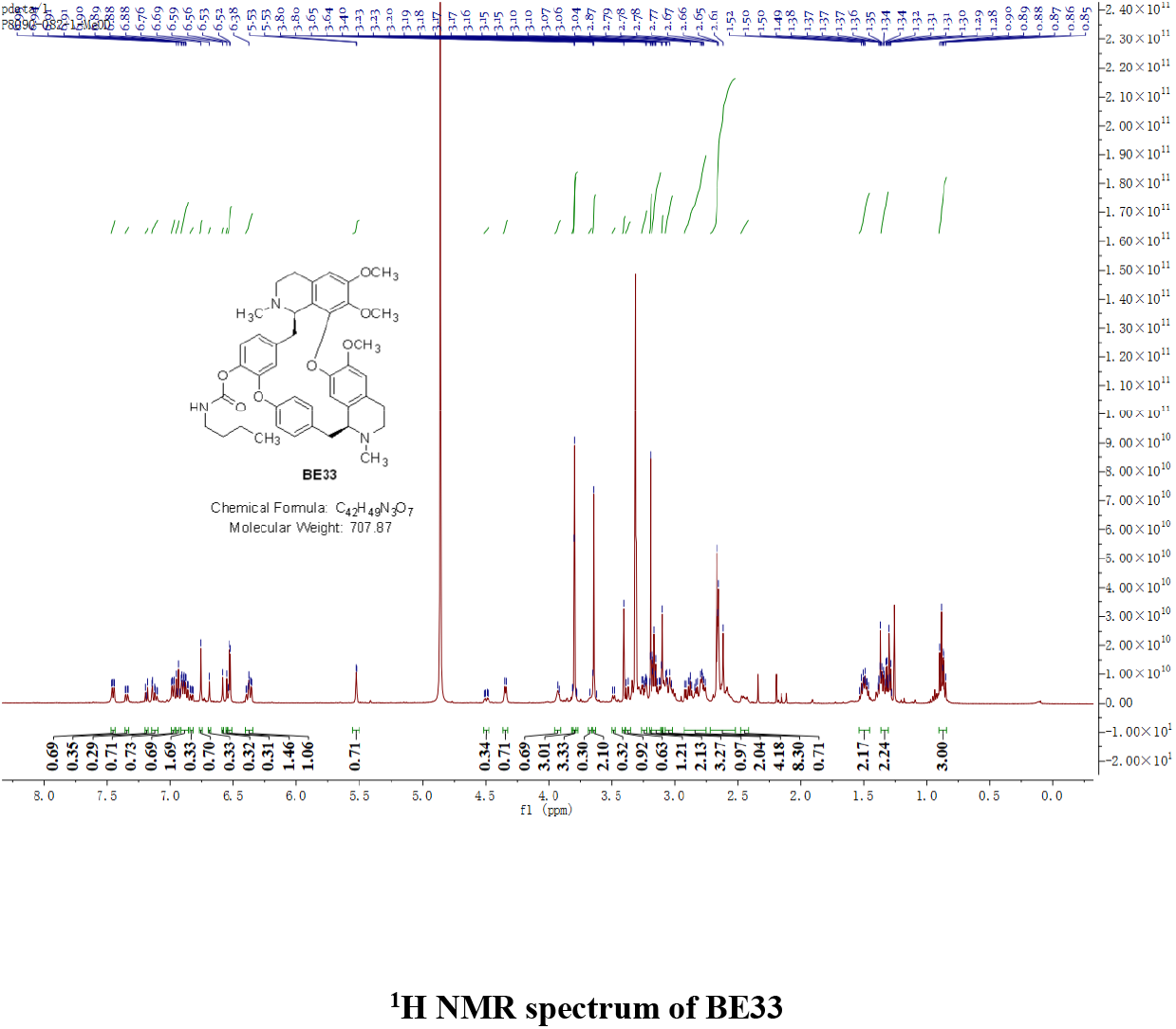
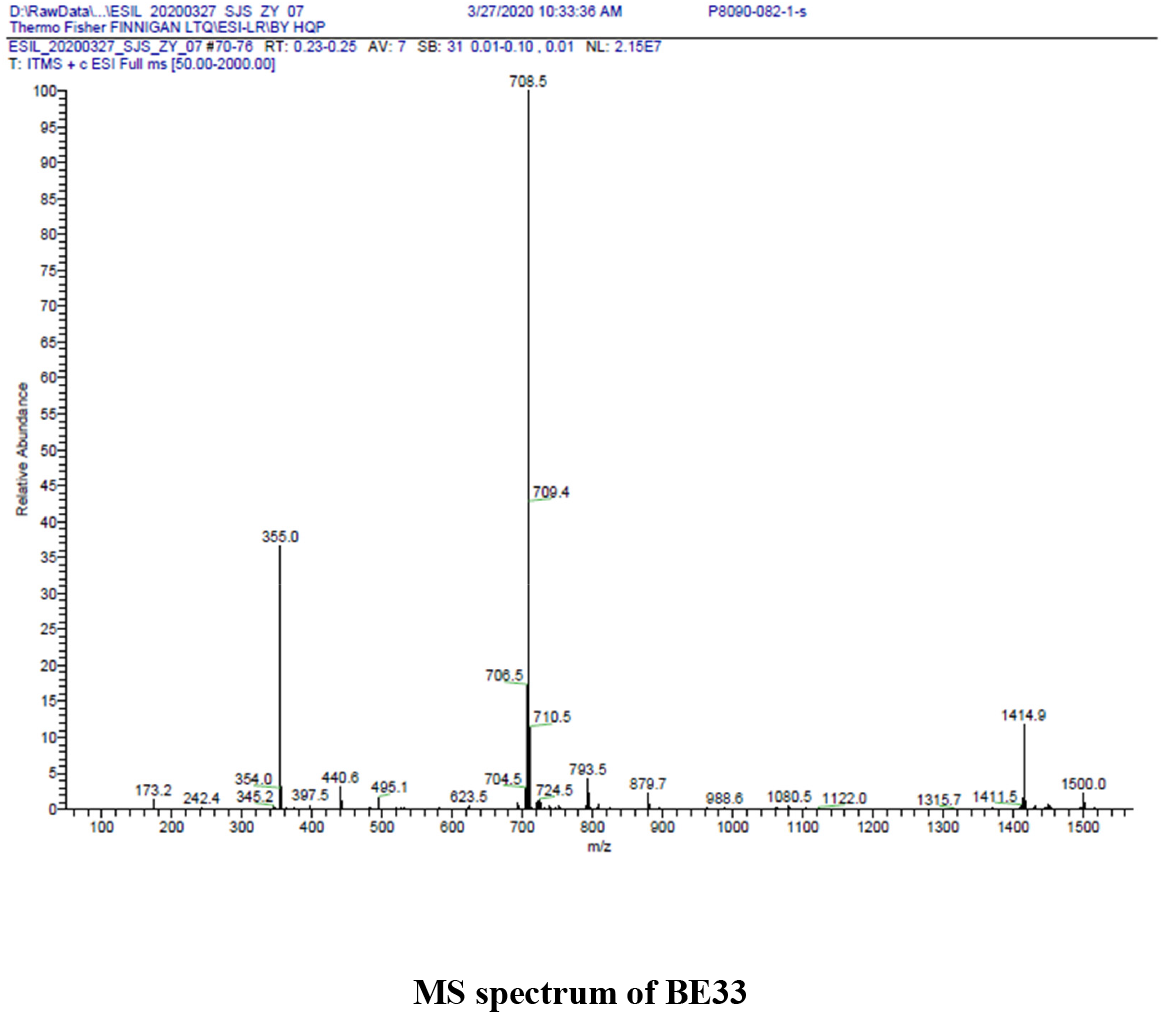
Spectral data for compounds BE12-1, BE30, BE31, BE32 and BE33. ^1^H NMR and MS spectra

## References and Notes

Banerjee, I., Behl, B., Mendonca, M., Shrivastava, G., Russo, A.J., Menoret, A., Ghosh, A., Vella, A.T., Vanaja, S.K., Sarkar, S.N., et al. (2018). Gasdermin D Restrains Type I Interferon Response to Cytosolic DNA by Disrupting Ionic Homeostasis. Immunity 49, 413–426.e415.

Bao, L., Deng, W., Huang, B., Gao, H., Liu, J., Ren, L., Wei, Q., Yu, P., Xu, Y., Qi, F., et al. (2020). The pathogenicity of SARS-CoV-2 in hACE2 transgenic mice.

Blanco-Melo, D., Nilsson-Payant, B.E., Liu, W.C., Uhl, S., Hoagland, D., Møller, R., Jordan, T.X., Oishi, K., Panis, M., Sachs, D., et al. (2020). Imbalanced Host Response to SARS-CoV-2 Drives Development of COVID-19. Cell 181, 1036–1045.e1039.

C., H., J. F.-W. C, Terrence Tsz-Tai Yuen, Huiping Shuai, Shuofeng Yuan, Yixin Wang, Bingjie Hu, Cyril Chik-Yan Yip, Jessica Oi-Ling Tsang, Xiner Huang, et al. (2020). Comparative tropism, replication kinetics, and cell damage profiling of SARS-CoV-2 and SARS-CoV with implications for clinical manifestations, transmissibility, and laboratory studies of COVID-19: an observational study. Lancet Microbe.

Cady, S.D., Schmidt-Rohr, K., Wang, J., Soto, C.S., DeGrado, W.F., and Hong, M. (2010). Structure of the amantadine binding site of influenza M2 proton channels in lipid bilayers. Nature 463, 689–U127.

Cao, X. (2020). COVID-19: immunopathology and its implications for therapy. Nat Rev Immunol 20, 269–270.

Castano-Rodriguez, C., Honrubia, J.M., Gutierrez-Alvarez, J., DeDiego, M.L., Nieto-Torres, J.L., Jimenez-Guardeno, J.M., Regla-Nava, J.A., Fernandez-Delgado, R., Verdia-Baguena, C., Queralt-Martin, M., et al. (2018). Role of Severe Acute Respiratory Syndrome Coronavirus Viroporins E, 3a, and 8a in Replication and Pathogenesis. Mbio 9.

Chen, N., Zhou, M., Dong, X., Qu, J., Gong, F., Han, Y., Qiu, Y., Wang, J., Liu, Y., Wei, Y., et al. (2020). Epidemiological and clinical characteristics of 99 cases of 2019 novel coronavirus pneumonia in Wuhan, China: a descriptive study. Lancet 395, 507–513.

Cohen, J.R., Lin, L.D., and Machamer, C.E. (2011). Identification of a Golgi complex-targeting signal in the cytoplasmic tail of the severe acute respiratory syndrome coronavirus envelope protein. J Virol 85, 5794–5803.

Conos, S.A., Chen, K.W.W., De Nardo, D., Hara, H., Whitehead, L., Nunez, G., Masters, S.L., Murphy, J.M., Schroder, K., Vaux, D.L., et al. (2017). Active MLKL triggers the NLRP3 inflammasome in a cell-intrinsic manner. P Natl Acad Sci USA 114, E961–E969.

de Wit, E., van Doremalen, N., Falzarano, D., and Munster, V.J. (2016). SARS and MERS: recent insights into emerging coronaviruses. Nat Rev Microbiol 14, 523–534.

Ding, J., Wang, K., Liu, W., She, Y., Sun, Q., Shi, J., Sun, H., Wang, D.C., and Shao, F. (2016). Pore-forming activity and structural autoinhibition of the gasdermin family. Nature 535, 111–116.

Feltham, R., and Vince, J.E. (2018). Ion Man: GSDMD Punches Pores to Knock Out cGAS. Immunity 49, 379–381.

Gong, Y.N., Guy, C., Olauson, H., Becker, J.U., Yang, M., Fitzgerald, P., Linkermann, A., and Green, D.R. (2017). ESCRT-III Acts Downstream of MLKL to Regulate Necroptotic Cell Death and Its Consequences. Cell 169, 286–300 e216.

Grifoni, A., Sidney, J., Zhang, Y., Scheuermann, R.H., Peters, B., and Sette, A. (2020). A Sequence Homology and Bioinformatic Approach Can Predict Candidate Targets for Immune Responses to SARS-CoV-2. Cell Host Microbe 27, 671–680.e672.

Guan, W.J., Ni, Z.Y., Hu, Y., Liang, W.H., Ou, C.Q., He, J.X., Liu, L., Shan, H., Lei, C.L., Hui, D.S.C., et al. (2020). Clinical Characteristics of Coronavirus Disease 2019 in China. N Engl J Med 382, 1708–1720.

Liao, Y., Yuan, Q., Torres, J., Tam, J.P., and Liu, D.X. (2006). Biochemical and functional characterization of the membrane association and membrane permeabilizing activity of the severe acute respiratory syndrome coronavirus envelope protein. Virology 349, 264–275.

Lippi, G., and South, A.M. (2020). Electrolyte imbalances in patients with severe coronavirus disease 2019 (COVID-19). Ann Clin Biochem 57, 262–265.

Louhaichi, S., Allouche, A., Baili, H., Jrad, S., Radhouani, A., Greb, D., Akrout, I., Ammar, J., Hamdi, B., Added, F., et al. (2020). Features of patients with 2019 novel coronavirus admitted in a pneumology department: The first retrospective Tunisian case series. Tunis Med 98, 261–265.

Mehta, P., McAuley, D.F., Brown, M., Sanchez, E., Tattersall, R.S., and Manson, J.J. (2020). COVID-19: consider cytokine storm syndromes and immunosuppression. Lancet 395, 1033–1034.

Merad, M., and Martin, J.C. (2020). Pathological inflammation in patients with COVID-19: a key role for monocytes and macrophages. Nat Rev Immunol 20, 355–362.

Nieto-Torres, J.L., DeDiego, M.L., Verdiá-Báguena, C., Jimenez-Guardeño, J.M., Regla-Nava, J.A., Fernandez-Delgado, R., Castaño-Rodriguez, C., Alcaraz, A., Torres, J., Aguilella, V.M., et al. (2014). Severe acute respiratory syndrome coronavirus envelope protein ion channel activity promotes virus fitness and pathogenesis. Plos Pathog 10, e1004077.

Nieto-Torres, J.L., Verdiá-Báguena, C., Jimenez-Guardeño, J.M., Regla-Nava, J.A., Castaño-Rodriguez, C., Fernandez-Delgado, R., Torres, J., Aguilella, V.M., and Enjuanes, L. (2015). Severe acute respiratory syndrome coronavirus E protein transports calcium ions and activates the NLRP3 inflammasome. Virology 485, 330–339.

Noris, M., Benigni, A., and Remuzzi, G. (2020). The case of Complement activation in COVID-19 multiorgan impact. Kidney Int.

Peeples, L. (2020). News Feature: Avoiding pitfalls in the pursuit of a COVID-19 vaccine. Proc Natl Acad Sci U S A 117, 8218–8221.

Pervushin, K., Tan, E., Parthasarathy, K., Lin, X., Jiang, F.L., Yu, D., Vararattanavech, A., Soong, T.W., Liu, D.X., and Torres, J. (2009). Structure and inhibition of the SARS coronavirus envelope protein ion channel. Plos Pathog 5, e1000511.

Surya, W., Li, Y., Verdia-Baguena, C., Aguilella, V.M., and Torres, J. (2015). MERS coronavirus envelope protein has a single transmembrane domain that forms pentameric ion channels. Virus Res 201, 61–66.

Tay, M.Z., Poh, C.M., and Rénia, L. (2020). The trinity of COVID-19: immunity, inflammation and intervention. Immunity 20, 363–374.

Verdiá-Báguena, C., Nieto-Torres, J.L., Alcaraz, A., Dediego, M.L., Enjuanes, L., and Aguilella, V.M. (2013). Analysis of SARS-CoV E protein ion channel activity by tuning the protein and lipid charge. Biochim Biophys Acta 1828, 2026–2031.

Wang, Y.P., Gao, W.Q., Shi, X.Y., Ding, J.J., Liu, W., He, H.B., Wang, K., and Shao, F. (2017). Chemotherapy drugs induce pyroptosis through caspase-3 cleavage of a gasdermin. Nature 547, 99–+.

Westerbeck, J.W., and Machamer, C.E. (2019). The Infectious Bronchitis Coronavirus Envelope Protein Alters Golgi pH To Protect the Spike Protein and Promote the Release of Infectious Virus. J Virol 93.

Wrapp, D., and Wang, N. (2020). Cryo-EM structure of the 2019-nCoV spike in the prefusion conformation. Science 367, 1260–1263.

Wu, Y., Guo, C., Tang, L., Hong, Z., Zhou, J., Dong, X., Yin, H., Xiao, Q., Tang, Y., Qu, X., et al. (2020). Prolonged presence of SARS-CoV-2 viral RNA in faecal samples. Lancet Gastroenterol Hepatol 5, 434–435.

Xia, B.Q., Fang, S., Chen, X.Q., Hu, H., Chen, P.Y., Wang, H.Y., and Gao, Z.B. (2016). MLKL forms cation channels. Cell Res 26, 517–528.

Xu, Z., Shi, L., Wang, Y., Zhang, J., Huang, L., Zhang, C., Liu, S., Zhao, P., Liu, H., Zhu, L., et al. (2020). Pathological findings of COVID-19 associated with acute respiratory distress syndrome. Lancet Respir Med 8, 420–422.

Zaki, A.M., van Boheemen, S., Bestebroer, T.M., Osterhaus, A.D., and Fouchier, R.A. (2012). Isolation of a novel coronavirus from a man with pneumonia in Saudi Arabia. N Engl J Med 367, 1814–1820.

Zhang, H., Zhou, P., Wei, Y., Yue, H., Wang, Y., Hu, M., Zhang, S., Cao, T., Yang, C., Li, M., et al. (2020). Histopathologic Changes and SARS-CoV-2 Immunostaining in the Lung of a Patient With COVID-19. Ann Intern Med 172, 629–632.

Zhong, N.S., Zheng, B.J., Li, Y.M., Poon, Xie, Z.H., Chan, K.H., Li, P.H., Tan, S.Y., Chang, Q., Xie, J.P., et al. (2003). Epidemiology and cause of severe acute respiratory syndrome (SARS) in Guangdong, People’s Republic of China, in February, 2003. Lancet 362, 1353–1358.

Zhu, N., Zhang, D., Wang, W., Li, X., Yang, B., Song, J., Zhao, X., Huang, B., Shi, W., Lu, R., et al. (2020). A Novel Coronavirus from Patients with Pneumonia in China, 2019. N Engl J Med 382, 727–733.

## References

Wang, D., Zheng, J., Hu, Q., Zhao, C., Chen, Q., Shi, P., Chen, Q., Zou, Y., Zou, D., Liu, Q., et al. (2020). Magnesium protects against sepsis by blocking gasdermin D N-terminal-induced pyroptosis. Cell Death Differ 27, 466–481.

